# Cell type-specific histone acetylation landscape in Alzheimer’s disease reveals a putative role of MITF in microglia

**DOI:** 10.64898/2026.09.09.750356

**Authors:** Charbel Gergian, Paulina Urbanaviciute, Philippa M. Wells, Janis L. Transfeld, Aydan Askarova, Reuben M. Yaa, Yukyee Wu, Kevin Chris Ziegler, Christian K. Nickl, Martina P. Pasillas, Johannes C. M. Schlachetzki, Inge R. Holtman, Peter T. Nelson, Robert A. Rissman, James B. Brewer, Sarah J. Marzi, Christopher K. Glass, Alexi Nott

**Affiliations:** Department of Brain Sciences, Imperial College London, London, UK; UK Dementia Research Institute, Imperial College London, London, UK; Department of Basic and Clinical Neuroscience, Institute of Psychiatry, Psychology and Neuroscience, King’s College London, London, UK; UK Dementia Research Institute, King’s College London, London, UK; Department of Cellular and Molecular Medicine, University of California San Diego, La Jolla, USA; Department of Neurosciences, University of California, San Diego, La Jolla, CA 92093, USA; Department of Biomedical Sciences, University Medical Center Groningen, University of Groningen, Groningen, The Netherlands; Sanders-Brown Center on Aging, University of Kentucky, Lexington, KY 40536, USA; Department of Physiology and Neuroscience, Alzheimer’s Therapeutic Research Institute, Keck School of Medicine of the University of Southern California, San Diego, CA; Department of Medicine, University of California San Diego, La Jolla, CA, USA

## Abstract

Alzheimer’s disease (AD) is characterised by aberrant amyloid beta and tau aggregation, neuroinflammation, demyelination and neurodegeneration, which have been linked to changes in cell-specific gene expression signatures. Among the mechanisms driving cell-type-specific transcriptional changes, histone acetylation plays a central role in regulating gene activity. While global alterations in histone acetylation have been implicated in AD, the contribution of individual cell types to these epigenetic changes remains poorly understood. To decode cell-type-specific changes in gene regulation in AD pathogenesis, we profiled histone H3 lysine 27 acetylation (H3K27ac) in microglia, oligodendrocytes and neurons from the prefrontal cortex of individuals with late-stage AD and non-dementia controls. Oligodendrocytes had the highest number of differential H3K27ac regions in AD, followed by microglia. Genes nearest to differential H3K27ac in purified microglia were enriched for phagocytosis, lipid processing, inflammatory and disease-associated cell state signature genes. Gene network analysis revealed downregulation of homeostatic genes in AD microglia and upregulation of immune activation, including signatures of lipid-handling and monocyte-derived macrophages. Oligodendrocyte co-regulated regions were indicative of increased MHC class I antigen presentation and altered neuron-oligodendrocyte interactions in AD. We identified H3K27ac allele-specific variants (ASVs) enriched near endolysosomal and ubiquitin-proteasome-associated genes in microglia and neurons. ASVs coincided with AD genome-wide association study (GWAS) risk loci, including *CLU* in oligodendrocytes and *HLA-DRB1* in microglia. DNA motif analysis identified putative transcription factor drivers of AD glial dysregulation, including the lysosomal-associated MITF, Cap’n’collar (CNC) family (BACH1 and NFE2) and AP-1 activation in microglia. DNA binding of the MITF protein in human microglia was localised to lysosomal-associated genes and enriched in H3K27ac regions upregulated in AD and near disease-associated microglia (DAM) genes. Collectively, these findings implicate lysosomal dysfunction and upstream transcriptional regulation via MITF as key processes in AD microglia.

## Introduction

AD is a complex polygenic condition characterised by progressive neurodegeneration, impaired synaptic connectivity, demyelination and neuroinflammation. Recent single-cell gene expression studies have identified underlying transcriptional changes in neuronal subtypes, oligodendrocytes and microglia^1-4^. These include selective vulnerability of excitatory and inhibitory neurons^1^, changes in myelination-related processes^2^, and shifts from homeostatic microglia to inflammatory and lipid-processing states^5,6^. Gene expression programs are orchestrated at the level of the epigenome, which integrates environmental and disease signals to elicit cellular responses^7^. Transcription factors mediate gene regulation and are key to understanding the molecular mechanisms underlying disease pathogenesis^8^. However, the epigenetic alterations and transcription factors that mediate AD-associated changes in cell-type gene expression are unclear.

The epigenetic modification histone H3 lysine 27 acetylation (H3K27ac) annotates gene regulatory regions, including promoters and distal enhancers^9^, and is dysregulated in the cortex of the AD brain^10,11^. H3K27ac profiling of cell-type-enriched nuclei for the major cell types of the brain showed that differential acetylation associated with amyloid beta load was most prevalent in oligodendrocytes, including at gene regulatory regions near genome-wide association study (GWAS) risk loci associated with late-onset AD^12^. In non-dementia conditions, AD GWAS variants are located near genes expressed in microglia^13-15^, and are enriched at enhancers in macrophages and microglia^16-20^. In addition, the myeloid lineage-determining transcription factor, PU.1, has been identified as a key putative regulator of AD risk^21,22^. However, cell type-specific gene regulatory regions associated with GWAS loci under AD conditions have been underexplored.

To understand how AD risk variants and epigenetic dysregulation intersect at cell-type-specific regulatory elements, we profiled H3K27ac in microglia, oligodendrocytes, and neurons from the prefrontal cortex of individuals with AD using chromatin immunoprecipitation followed by sequencing (ChIP-seq). Dysregulated gene regulatory regions were identified as differential H3K27ac peaks near genes associated with immune responses, cell adhesion and endolysosomal trafficking in microglia, neuron-oligodendrocyte communication and immune signalling in oligodendrocytes, and protein trafficking and synaptic signalling in neurons. ASVs implicated common genetic determinants of gene regulation near endolysosomal and proteasomal-associated genes in microglia and coincided with AD GWAS loci. DNA-binding motif analysis and transcription factor profiling implicated a role for the lysosomal-associated microphthalmia-associated transcription factor (MITF) in AD-mediated microglial gene regulation.

## Results

### Identification of the cell-type regulatory landscapes

To assess changes in gene regulatory regions in microglia, oligodendrocytes and neurons in AD, we isolated cell-type-enriched nuclei populations from 15 non-dementia (ND) controls (Braak stage 0-II) and 26 individuals with late-stage AD (Braak stage V-VI) from the pre-frontal cortex (**Figure 1A and Table S1**). Cell-type-enriched nuclei were isolated using fluorescence-activated nuclei sorting (FANS)^16,23^ by immunostaining for the nuclear factors PU.1 (microglia), OLIG2 (oligodendrocytes) and NeuN (neurons) (**Figure 1A and 1B**). Genome-wide distribution of H3K27ac was mapped by ChIP-seq and used to annotate cell type gene regulatory regions, including promoters and enhancers (**Figure 1A**). In total, 94 H3K27ac samples (microglia 13 ND, 8 AD; oligodendrocytes 14 ND, 23 AD; neurons 15 ND, 21 AD) passed quality controls for the number of unique reads, fraction of reads in peaks (FRiP), and immunoprecipitation enrichment (normalized strand coefficient [NSC] and relative strand correlation [RSC]) (**Figure S1A-E**). Gene regulatory regions were defined for each cell type as H3K27ac peaks, and H3K27ac peak enrichment was determined for individual samples (**Table S2 and S3**). Cell type proportion of FANS-enriched nuclei was estimated by generating cell type-specific histone acetylation profiles for each sample against a reference human dataset of resected cortical samples^16^. Samples with an estimated high purity of the target cell type were retained, and the predicted cell type proportions were comparable for ND controls and AD (**Figure 1C and S2A**). H3K27ac samples clustered according to the cell type of origin by principal component analysis and Pearson’s correlation (**Figures S2B and S2C**). To assess nuclei enrichment by FANS, cell type-specific promoters were defined as gene regulatory regions proximal to transcriptional start sites (TSSs), followed by differential H3K27ac between cell types. Cell-type-specific promoters were enriched for signature genes of the corresponding cell type defined by human cortical single-nuclei (sn)RNA-seq^1^ using Expression Weighted Celltype Enrichment (EWCE) and correlation-adjusted mean rank (CAMERA) competitive gene set testing (**Figures 1D and 1E**). Cell-type-specific promoters for PU.1 nuclei were enriched for microglia ontology terms, OLIG2 nuclei were enriched for oligodendrocytes and oligodendrocyte precursor cells (OPCs), and NeuN nuclei for neuronal subtypes (**Figure 1F**). Collectively, FANS of postmortem cortical samples followed by H3K27ac ChIP-seq defined the gene regulatory landscape of microglia, oligodendrocytes and neurons in AD as well as ND controls.

**Figure 1.**
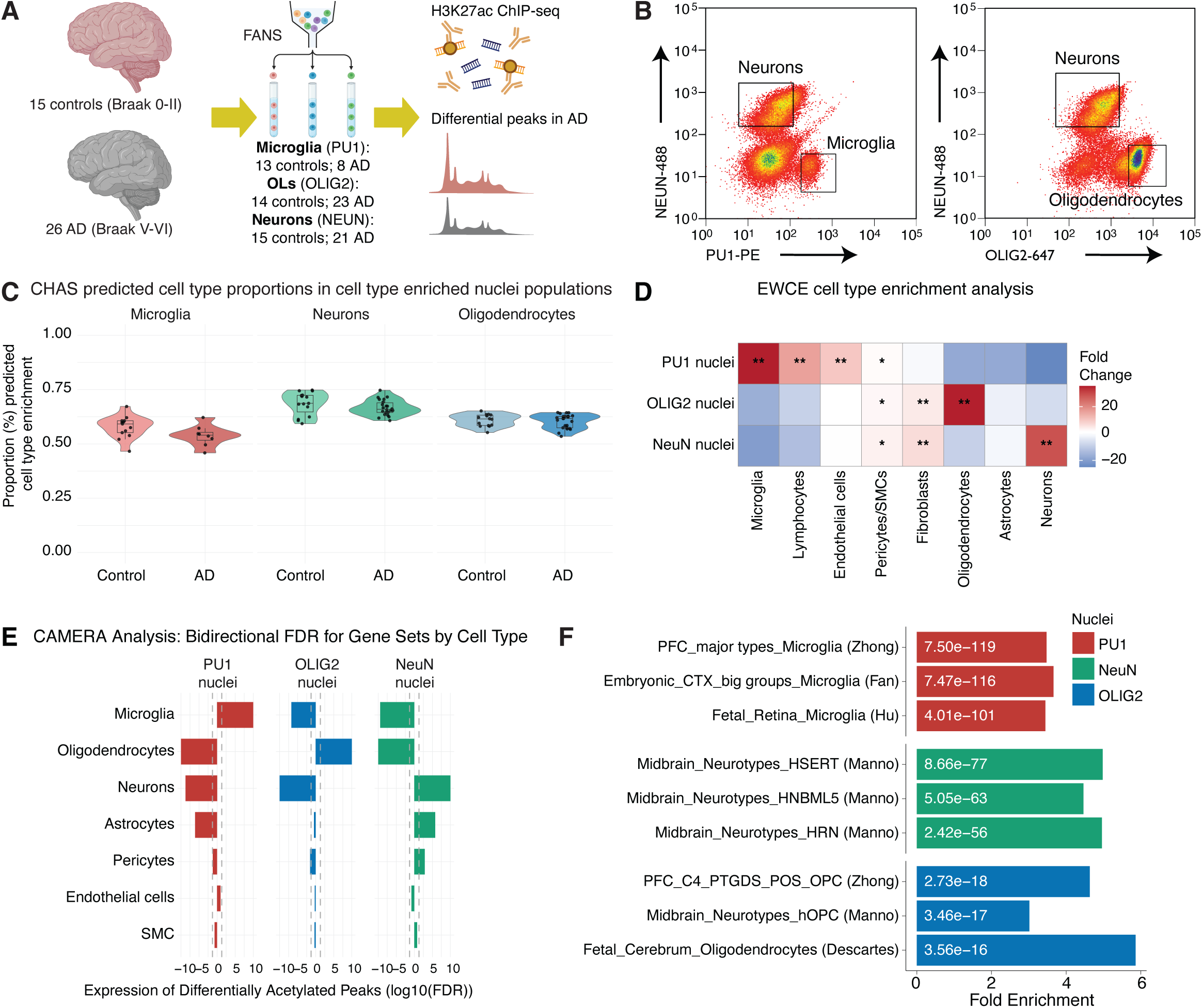
The epigenome of oligodendrocytes, neurons and microglia in AD. **(A)** Experimental design of PU.1^+^ (microglia/myeloid cells), OLIG2^+^ (oligodendrocyte) and NeuN^+^ (neuronal) nuclei from 15 control (Braak 0-II) and 26 AD (Braak V-VI) prefrontal cortex samples subjected to differential H3K27ac ChIP-seq analysis. **(B)** FANS gating strategy to enrich for microglia, oligodendrocyte and neuronal nuclei using the cell type nuclei markers PU.1, OLIG2 and NeuN, respectively. **(C)** Violin plots of the percentage proportion of the enriched cell types predicted using CHAS. **(D)** EWCE enrichment analysis of cell type-specific promoters in brain single-cell gene expression^80^. ^∗^Benjamini-Hochberg-corrected q < 0.05; ^∗∗^Benjamini-Hochberg-corrected q = 0. **(E)** CAMERA analysis for enrichment of brain cell type promoters against cell type marker genes defined by single-cell gene expression data^80^. Cell type-specific promoters were ranked relative to differential single-cell gene expression, accounting for inter-gene correlations. **(F)** Enrichment analysis of cell type-specific promoters for microglia, oligodendrocytes and neurons for cell type signature gene sets. Shown are the top 3 most significant terms per cell type.

### Differential histone acetylation of gene regulatory regions in AD

Gene regulatory regions that were differentially acetylated in AD compared to ND controls were identified by differential analysis of H3K27ac ChIP-seq peaks using DESeq2 (FDR < 0.05). The highest number of differential H3K27ac gene regulatory regions was identified in oligodendrocytes (1794 hyperacetylated; 2100 hypoacetylated), followed by microglia (1062 hyperacetylated; 1149 hypoacetylated), and neurons (196 hyperacetylated; 492 hypoacetylated) (**Figure 2A and Table S4**). The majority of differential H3K27ac gene regulatory regions were located within intronic and intergenic regions, indicating that in all three cell types, gene regulatory regions distal to TSSs were often affected in AD (**Figure S3A**). Functional enrichment analysis of the differential H3K27ac gene regulatory regions was assessed using gprofiler^24^ (**Table S5**). Gene regulatory regions that were hyperacetylated in microglia in AD were associated with immune and inflammatory responses, cell migration and adhesion, and vesicles including genes linked to lysosomal enzymes (*CTSD*, *CTSL*) endolysosome trafficking/positioning (*BORCS5*, *SNX10*, *SLC9A8*, *LRRK2* and *LGALS3*) and autophagy (*ATG7*) (**Figure 2B**), whereas microglia hypoacetylated regions in AD were associated with cell communication (**Figure S3B**). In oligodendrocytes, hyperacetylated regions were associated with cytoskeleton organisation, cell adhesion and neuronal processes (**Figure 2C**), and hypoacetylated regions were associated with metabolic processes and chromatin organisation (**Figure S3C**). Both hyperacetylated and hypoacetylated regions were nearby genes linked to neuronal processes, indicating potential dysregulated crosstalk with neurons (**Figure 2C and S3C**). H3K27ac regions upregulated in neurons were associated with protein localization and trafficking, receptor recycling and translational regulation (**Figure S3D**) and neuronal hypoacetylation was associated with dendrites and synapses, suggesting dysregulated synaptic function and neuronal signalling (**Figure S3E**).

**Figure 2.**
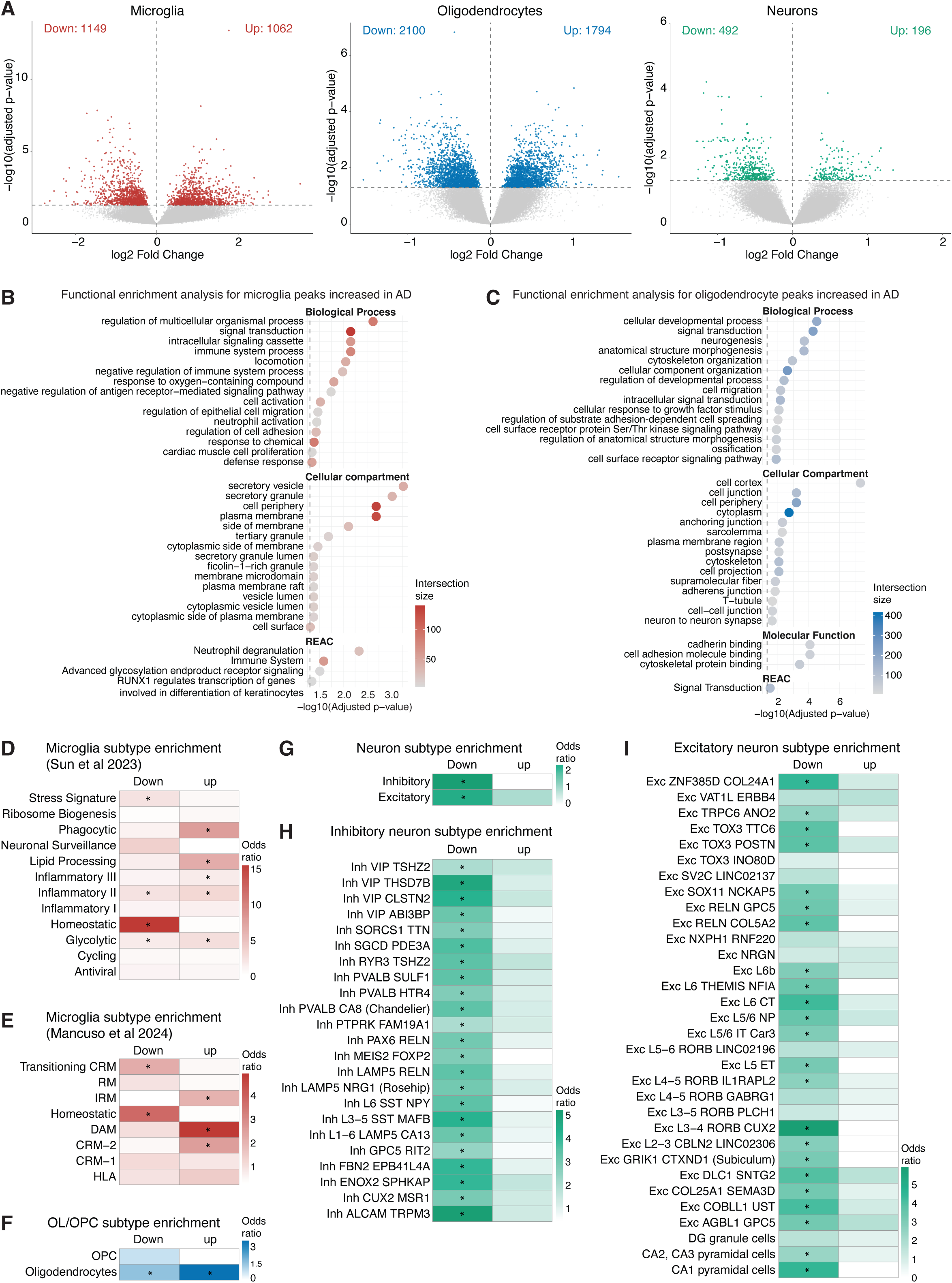
Cell-type epigenomic dysregulation in AD. **(A)** Volcano plots of DESeq2 differential H3K27ac analysis between AD and control samples in microglia, oligodendrocytes and neurons. **(B)** Functional enrichment analysis of annotated hyperacetylated regions in microglia using gprofiler. **(C)** Functional enrichment analysis of annotated hyperacetylated regions in oligodendrocytes using gprofiler. **(D)** Fisher’s exact test for enrichment of annotated microglial differential H3K27ac peaks for signature genes of microglial transcriptional substates defined using human postmortem brain snRNA-seq^4^. **(E)** Fisher’s exact test for enrichment of annotated microglial differential H3K27ac peaks for signature genes of microglial transcriptional substates defined using a human-mouse amyloid model by scRNA-seq^25^. **(F)** Fisher’s exact test for enrichment of annotated oligodendrocyte differential H3K27ac peaks for signature genes of oligodendrocytes and OPCs defined by snRNA-seq^3^. **(G-I)** Fisher’s exact test for enrichment of annotated neuronal differential H3K27ac peaks for signature genes of **(G)** excitatory and inhibitory neurons, **(H)** excitatory neuronal subtypes and **(I)** inhibitory neuronal subtypes defined by snRNA-seq^3^.

Dysregulated H3K27ac gene regulatory regions were assessed for enrichment of signature genes for cell subtypes and states using snRNA-seq datasets for microglia^4,25^ and cortical cell types^3^. Gene regulatory regions hypoacetylated in microglia in AD were strongly enriched for microglial homeostatic signature genes^4,25^ (**Figure 2D and 2E**), which is aligned with the downregulation of homeostatic genes in disease microglia^26,27^. In contrast, hyperacetylated regions were enriched for signature genes of phagocytic, lipid processing, and inflammatory transcriptional cell states defined using AD postmortem brains^4^. Consistent with this, hyperacetylated regions were associated with signature genes of DAM, interferon-response (IRM) and cytokine-response (CRM) microglial substates defined using a human-mouse amyloid model^25^ (**Figure 2D and 2E**), indicating a shift in AD from a homeostatic to an activated transcriptional state. Similar changes in microglial heterogeneity have been observed in single-nuclei gene expression studies for AD^4^. Dysregulated oligodendrocyte H3K27ac regions predominantly captured oligodendroglial signature genes^1^ (**Figure 2F**). Hypoacetylated regions in oligodendrocytes were also modestly enriched for OPC signature genes^1^, indicating that progenitor cells are underrepresented or dysregulated in AD (**Figure 2F**). Hypoacetylation in neurons was broadly enriched for both inhibitory and excitatory signature genes^1^ (**Figure 2G-I**), potentially reflecting changes in neuronal subtype proportions. Signatures of Sst, Pvalb, Vip, and Lamp5 interneurons were downregulated in AD and have previously been reported as vulnerable in AD by snRNA-seq^1,28^ (**Figure 2H**). Likewise, signatures of RORB-expressing and superficial-layer (L2/3 IT) excitatory neurons were downregulated in AD, consistent with reduced abundance of these populations reported in AD snRNA-seq datasets^1,28,29^ (**Figure 2I**). Similarly, signatures genes for TOX3^+^TTC6^+^, and RORB-positive L1 (RORB^+^GPC5^+^) and L5 (AGBL1^+^GPC5^+^) excitatory neurons were hypoacetylated (**Figure 2I**) and were shown to be vulnerable in AD using imaging mass cytometry^30,31^. Signatures associated with vulnerable neurons of the entorhinal cortex (RELN+) and hippocampus (CA1 pyramidal neurons) were also identified as downregulated in AD (**Figure 2I**), possibly reflecting shared transcriptional signatures across neuronal subtypes.

### Co-regulated gene regulatory regions in AD

To identify gene regulatory regions that are co-regulated across samples, we adapted weighted gene co-expression network analysis (WGCNA) for H3K27ac peaks (see methods). WGCNA identified 17, 14 and 10 modules of co-regulated gene regulatory regions for microglia, oligodendrocytes and neurons, respectively (**Figure 3A-C, S4 and Table S6**). Module trait analysis identified two microglial modules with a positive correlation with AD (M1 and M2) and two modules with a negative correlation (M16 and M17) (**Figure 3A**). Microglia M1 had the strongest positive correlation with AD (r = 0.78, p<0.001), with enrichment terms associated with cytokine activity and plasma membrane (**Figure 3D and Table S7**). Module M1 included genes characteristic of monocyte-derived macrophages (e.g. *MS4A* cluster, *SIGLEC1*, *CLEC5A*/*4E*, *FPR3*) and lymphocytes (*CD5*, *CD72*, *CD79B*, *FCER2*, *FCRLB*, *PDCD1*, *IL12RB2*), as well as lipid handling genes (*CD9*, *LGALS3BP*, *ABCA1*, *MGLL* and *UGCG*). The downregulated microglia module M16 was enriched for signalling pathways associated with regulation of immune processes and activation (**Figure 3E**). This module included canonical human homeostatic microglia genes (including *CX3CR1*, *P2RY13*, *TGFBR1*, *IRF8*, *SELPLG*, *BHLHE41*) of which *SALL1*, *MEF2C* and *IL6ST* were top-ranked hub genes (**Figure 3F and Table S6**), as well as 6 AD GWAS genes (*BIN1*, *PICALM*, *TREM2*, *BLNK*, *SCIMP* and *TSPAN14*). When comparing with published single-cell RNA-seq data^25^, microglia M1 genes were associated with signature genes of DAM and antigen-presenting response (HLA) microglia substates, and M16 had the strongest enrichment for the homeostatic substate (**Figure 3G**). M2 and M17 were not enriched for specific functional pathways. However, M2 genes were related to the proteasome (*PSMD2*), DNA repair (e.g. *NEIL1*, *NTHL1*), and metabolism (e.g. *PFKL*, *UQCRC1*, *BNIP3*, *ALDH7A1*, *HAGH*), while M17 included lysosomal and lipid-related genes (e.g. *NR1H3*, *VLDLR*, *LAPTM4B*, *GPR137B*) (**Table S6**).

**Figure 3.**
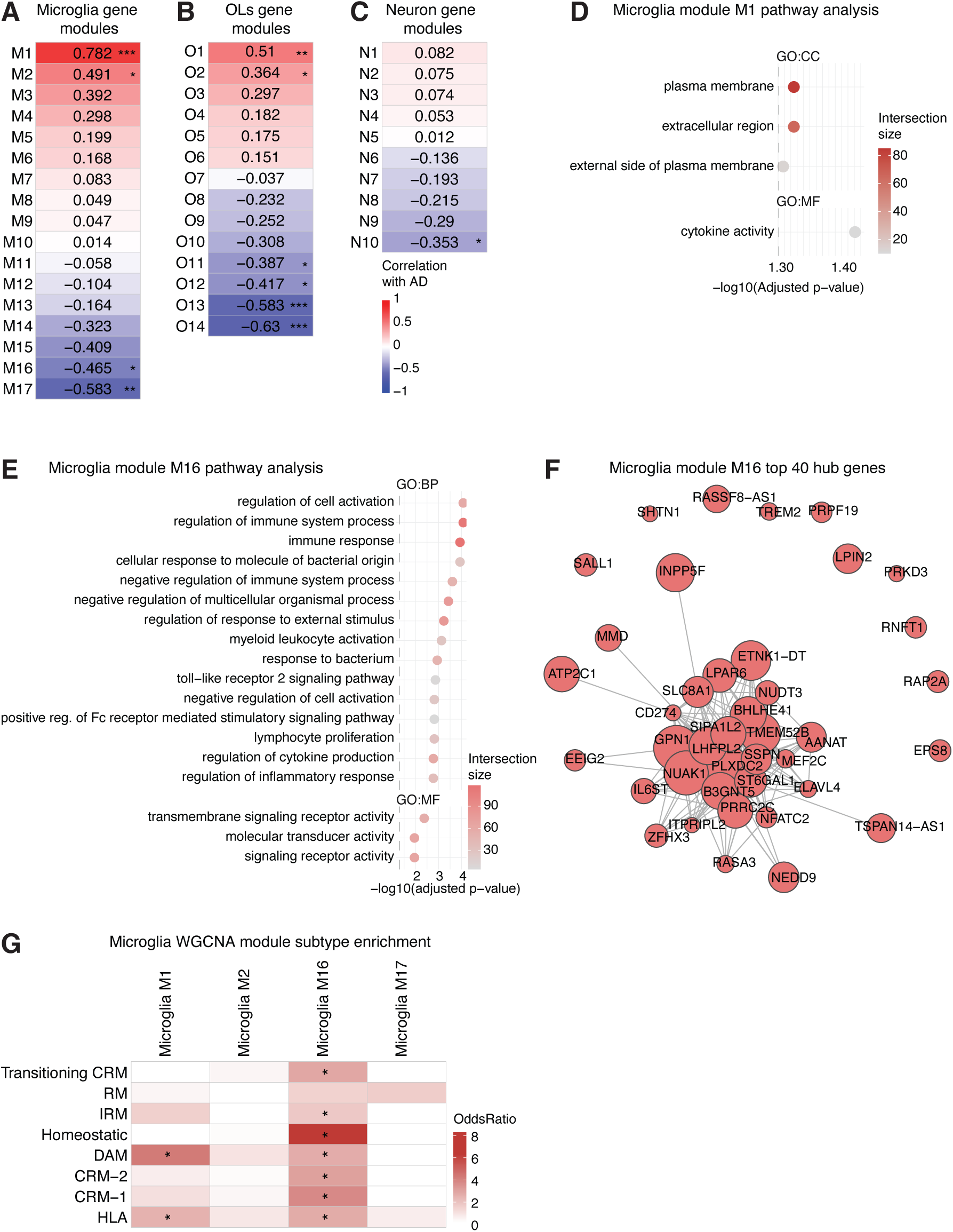
Weighted gene co-expression network analysis and cell subtype enrichment. **(A)** Heatmap of microglia module correlations with AD status. **(B)** Heatmap of oligodendrocyte module correlations with AD status. **(C)** Heatmap of neuron module correlations with AD status. **(D)** Functional enrichment analysis of microglia module M1 genes using gprofiler. **(E)** Functional enrichment analysis of microglia module M16 genes using gprofiler. **(F)** Network visualization of top 40 hub genes for microglia module M16. **(G)** Fisher’s exact test for enrichment of microglia module genes for signature genes of microglial transcriptional substates defined using a human-mouse amyloid model by scRNA-seq^25^. Correlation p values; * p<0.05, ** p < 0.01, *** p < 0.001.

Six oligodendrocyte modules were significantly correlated with AD, two modules with a positive correlation (O1 and O2) and four with a negative correlation (O11-O14) (**Figure 3B**). The upregulated oligodendrocyte O1 module was not enriched for any functional pathways. However, O1 hub genes were related to autophagy and lysosomal processes, including *SCYL1* and *CTSF,* suggesting broad lysosomal dysregulation of glia in AD (**Table S6**). Additionally, O1 included the AD risk gene *IQCK* (**Table S6**). The oligodendrocyte module O2 was enriched for functional pathways related to MHC class I antigen processing and presentation (*HLA-B*, *HLA-C*, *HLA-F*, *NLRC5, ISG15*, *ICAM1* and *IL6R*), suggesting acquisition of immune-reactive features characteristic of disease-associated oligodendrocytes (**Figure S5A and Table S7**). The downregulated oligodendrocyte O11 module was enriched for terms associated with development and differentiation, as well as chromatin and DNA binding-associated terms, providing further evidence of gene regulatory dysfunction in AD (**Figure S5B and Table S7**). The negatively correlated module O12 was enriched for terms associated with OPC identity and neuron-oligodendrocyte interface (**Figure S5C and Table S7**). This module included OPC markers (*PTPRZ1, SOX5, MYT1)* as well as extracellular matrix proteins (e.g. *VCAN, COL9A1, LAMA4),* adhesion and GPCR-signalling (**Table S6**), consistent with an oligodendrocyte program supporting neuronal and synaptic integrity. Modules O13 and O14 were not enriched for any functional pathways. However, O13 included the AD risk gene *APOE* and transcription factors *MITF*, *SALL1* and *MEF2C* (**Table S6**). The top 35 hub genes for O14 included subunits of the tau phosphatase (*PPP2R3A* and *PPP2R2C*) and *PLD3*, which has been implicated in late-onset AD risk^32^ (**Table S6**). Lastly, the neuronal N10 module had a significant negative correlation with AD (**Figure 3C**) and consisted mostly of noncoding lincRNA, LOC and miRNAs, and olfactory genes (**Figure S5D and Table S6**).

### Allele-specific histone acetylation signatures are linked to glial AD genetic susceptibility

AD genetic heritability has been associated with microglia and macrophage enhancers^16,18,19,21^. To determine whether gene regulatory regions defined using postmortem AD and ND controls are associated with AD heritability, we used stratified linkage disequilibrium score (LDSC) regression analysis. As expected, we found that AD heritability was associated with microglia and was not significant for oligodendrocytes and neurons (**Figure 4A**). Common variants linked to disease have been proposed to alter the function of gene regulatory regions. H3K27ac provides a proxy of enhancer and promoter activity and was used to perform allele-specific histone modification analysis of genome-wide common variants. SNP coverage, allelic balance and mapping bias were similarly distributed across the three cell types (**Figure S6A-D**). H3K27ac ASV analysis identified 353, 2326 and 921 loci with allele-specific histone acetylation in microglia, oligodendrocytes and neurons, respectively, which were mostly cell type-specific (**Figure 4B and Table S8**). The distribution of ASVs was most prominent at promoter peaks, followed by intronic and distal intergenic peaks (**Figure S6E**).

**Figure 4.**
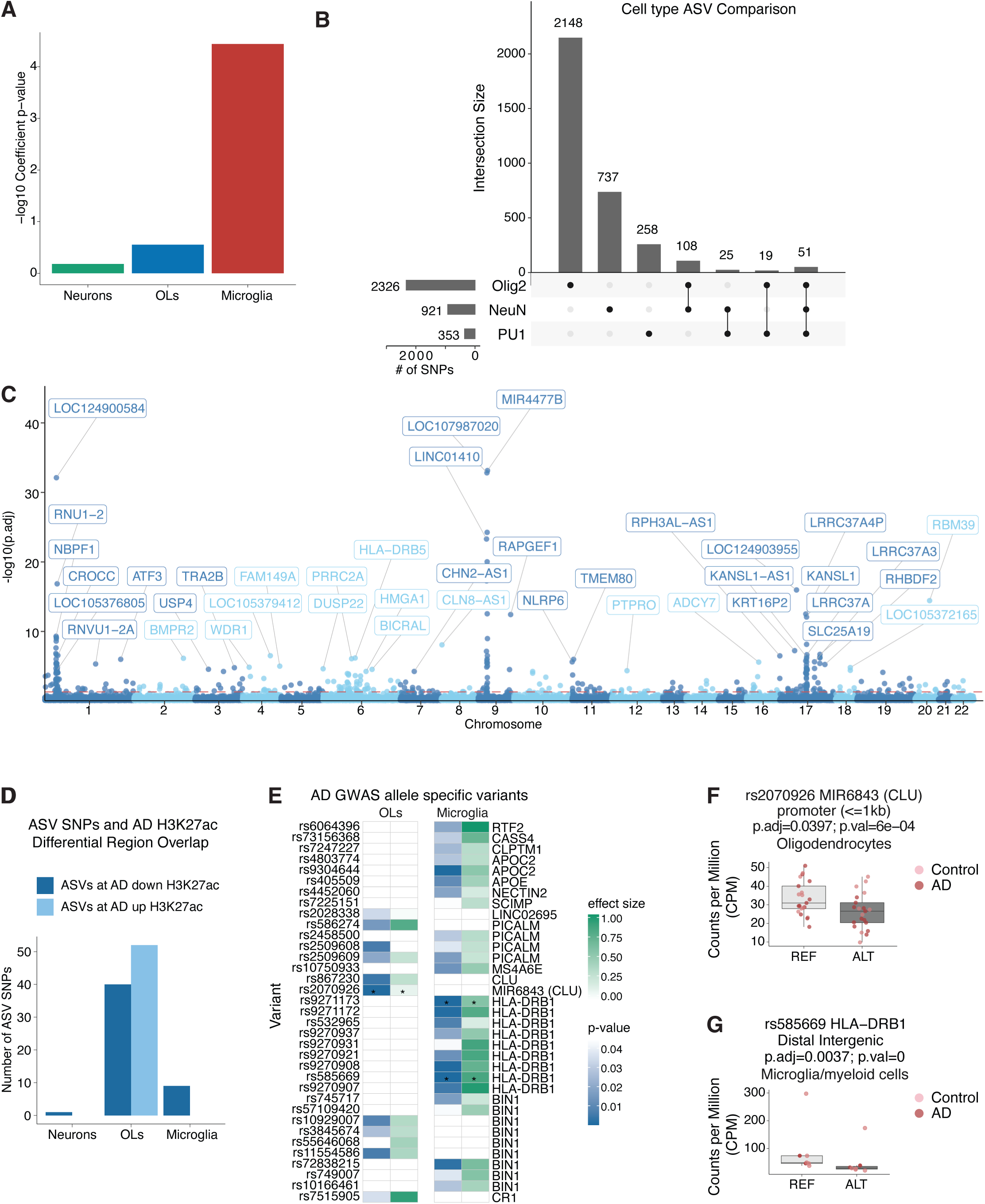
Genetic risk of AD is associated with microglia. **(A)** Stratified LDSC regression analysis using GWAS summary statistics for AD^15^ and H3K27ac regions identified in microglia, oligodendrocytes and neurons. **(B)** Upset plot of the number of unique and shared ASVs in microglia, oligodendrocytes, and neurons. **(C)** Manhattan plot of microglia ASVs with the top 40 annotated to proximal genes. **(D)** Number of ASVs overlapping H3K27ac regions that are differential in AD. **(E)** Nominally significant ASVs that were identified as significant AD GWAS risk variants^15^ in oligodendrocytes and microglia. Shown are allelic-specific variant p-values and major allele combined effect sizes; * FDR p-value<0.05. **(F-G)** Boxplots of allelic read counts, normalised for library size as counts per million (CPM), at FDR-significant SNPs rs2070926 at the *CLU* locus in oligodendrocytes and **(F)** rs585669 at the *HLA-DRB1* locus in microglia **(G)**. Boxplots indicate the median and interquartile range (IQR), with whiskers representing 1.5×IQR. Data points are coloured pale red for control and dark red for AD.

Microglia ASVs were linked to genes associated with lysosomes (*RAB5A*, *RAB20*, *RAB31*, *LGMN*, *RUBCN*, *WIPI2*)^33^, proteasomes (*UBE2D3*, *USP4*, *USP22*, *KLHL24*), stress response (*ATF3*), and immune response (*IL4R*, *HLA-DQB1*), as well as cell identity (*SALL1*) and survival (*CSF1R*) (**Figure 4C and Table S8**). Microglial ASVs were also identified near genes linked to AD pathology (*SPP1*)^34^, familial dementia (*ITM2B*)^35^ and frontotemporal dementia and amyotrophic lateral sclerosis (*C9orf72*)^36^ (**Table S8**). Oligodendrocyte ASVs were associated with cell identity genes (*MOBP*, *MYRF*), including MBP, which binds to amyloid beta and is a biomarker of ageing and neurodegeneration^37^ (**Figure S6F and Table S8**). Interestingly, neuronal ASVs were linked to genes associated with processes identified for microglia ASVs, including the endosomal-lysosomal and ubiquitin-proteasome system (*RAB7A*, *RAB5B*, *RAB6B*, *RAB6C*, DNAJ (HSP40) family, HSP90 family, *PSMA4*), as well as familial dementia (*ITM2B*) (**Figure S6G and Table S8**).

To test whether genetics may contribute to differential H3K27ac in AD, ASVs were overlapped with AD differential H3K27ac regions. The number of ASVs overlapping differential H3K27ac regions was nine, 92, and one, for microglia, oligodendrocytes and neurons, respectively (**Figure 4D**). This suggests that in oligodendrocytes, genetics may contribute to altered H3K27ac in AD; however, this may reflect in part the lower number of differential H3K27ac regions identified in the other cell types. To assess whether allele-specific histone acetylation was associated with genetic susceptibility for AD, ASVs were subsetted to AD GWAS significant variants (p<5e-8)^15^. AD GWAS significant variants were identified as ASVs proximal to *MIR6843* at the *CLU* locus in oligodendrocytes (rs2070926) and at the *HLA-DRB1* locus in microglia (rs9271173 and rs585669) (**Figure 4E-G and S7A**). In addition, 26 and 11 nominally significant ASVs were identified as GWAS-significant variants in microglia and oligodendrocytes, respectively (**Figure 4E and S7A-B**). A second oligodendrocyte ASV was identified at the *CLU* locus (rs867230) (**Figure 4E and S7B**), and multiple ASVs were identified at the *BIN1* and *PICALM* loci in microglia and oligodendrocytes (**Figure 4E and S7A-B**). Multiple microglia nominal ASVs were identified at the *HLA-DRB1* and *APOE* loci (including *NECTIN2* and *APOC2*), as well as single ASVs at *CASS4* (rs73156368) and *MS4A* loci (rs10750933) (**Figure 4E, 4G and S7A**). Collectively, these findings implicate a contribution of AD genetic risk to the function of gene regulatory regions in microglia and oligodendrocytes.

### DNA motif analysis identified putative transcription factor drivers of AD glial dysregulation

Gene regulation is orchestrated through the recruitment and binding of transcription factors at enhancers and promoters that establish cell-type expression programmes. Transcription factors active in each cell type were inferred using promoter histone acetylation at the TSS of transcription factor genes^38^. Differential H3K27ac analysis identified 36 hyperacetylated regions proximal to 25 transcription factor genes in AD microglia, including *CEBPG*, *MITF*, *MAFB*, *RARA*, and *RUNX3*, and 21 hypoacetylated regions proximal to 16 transcription factor genes, including *FOXK1*, *RUNX2*, *CREB1*, and *TCF7L2* (**Figure 5A**). Several transcription factor families showed divergent regulation (e.g., RUNX3 up/RUNX2 down, NFATC2 up/NFATC1 down, IRF4 up/IRF2 down), which may indicate a functional shift between family members (**Figure 5A**). Hypoacetylated regions in AD oligodendrocytes were proximal to regulators of differentiation (*SOX1*, *SOX4*, *SOX11*, *TCF7L1*) and hyperacetylated at the maturation factor *CREB3L1* (**Figure 5A**), indicating altered OPC-to-oligodendrocyte programmes^39^. The liver X receptor alpha (LXRα) gene, *NR1H3*, was hypoacetylated in AD oligodendrocytes, indicating disrupted myelin lipid metabolism. *MITF* proximal-peaks were hyper and hypoacetylated in oligodendrocytes (**Figure 5A**), pointing to a broader dysregulated glial autolysosomal response in disease. In neurons, the ER stress/unfolded protein response factors *ATF6* and *CREB3L1* were downregulated in AD (**Figure 5A**), consistent with known disruption of ER proteostasis in neurodegeneration^40^.

**Figure 5.**
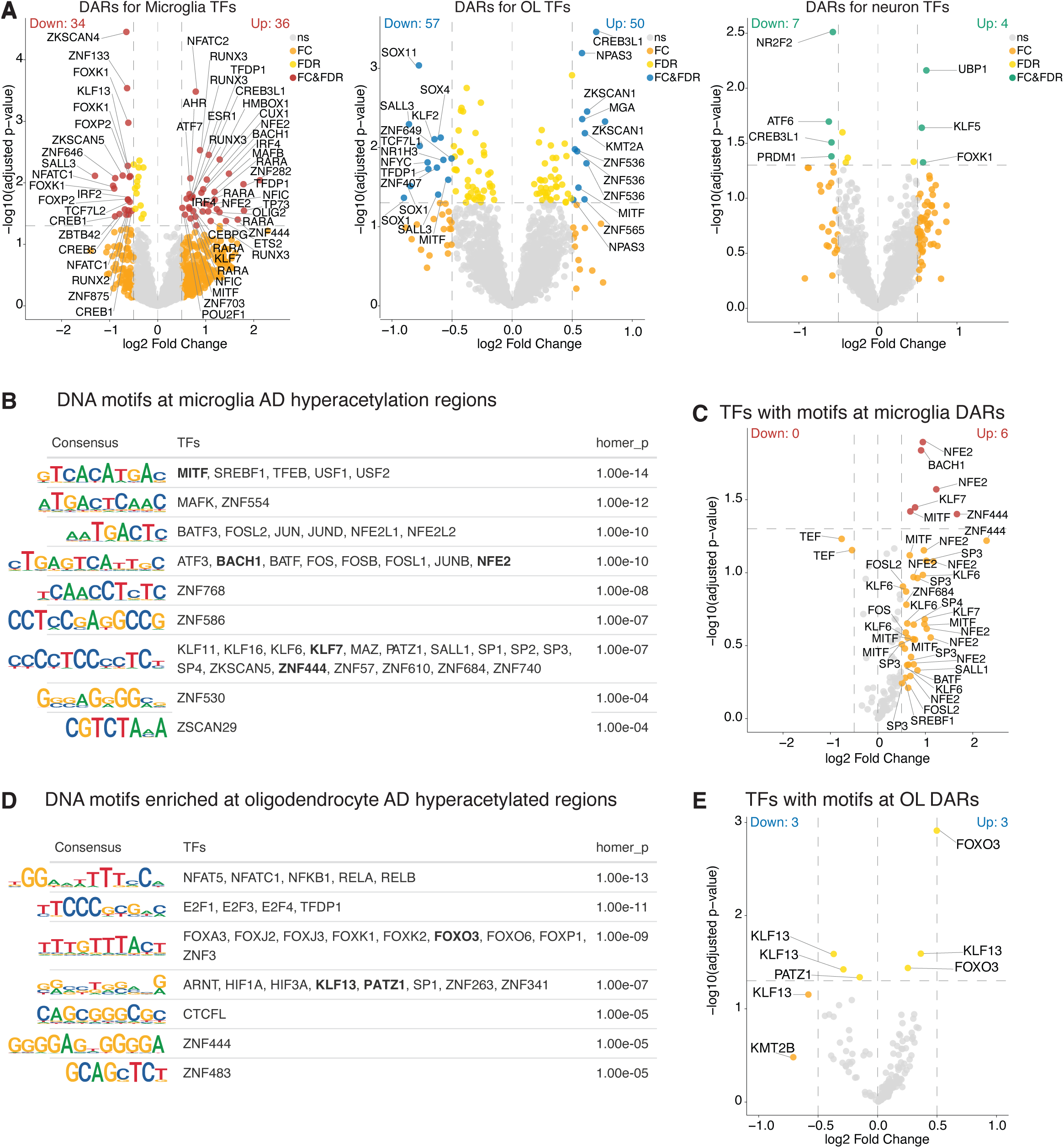
DNA motif analysis identified putative transcription factor drivers of AD dysregulation in microglia and oligodendrocytes. **(A)** Volcano plot of differential H3K27ac regions annotated to transcription factors in microglia, oligodendrocytes and neurons. (B) *De novo* motif analysis of microglia H3K27ac regions hyperacetylated in AD. *De novo* motifs were matched to transcription factors using Tomtom and filtered on transcription factors with active promoters in microglia. Motifs in bold are motifs matched to transcription factors with proximal differentially acetylated regions in AD. **(C)** Volcano plot of differential H3K27ac regions annotated to transcription factors with motifs enriched at microglia H3K27ac regions hyperacetylated and hypoacetylated in AD. (D) *De novo* motif analysis of oligodendrocyte H3K27ac regions hyperacetylated in AD. *De novo* motifs were matched to transcription factors using Tomtom and filtered on transcription factors with active promoters in oligodendrocytes. Motifs in bold are motifs matched to transcription factors with proximal differentially acetylated regions in AD. **(E)** Volcano plot of differential H3K27ac regions annotated to transcription factors with motifs enriched at oligodendrocyte H3K27ac regions hyperacetylated and hypoacetylated in AD.

A hypergeometric optimization of motif enrichment (HOMER) analysis was performed to determine whether differential H3K27ac regions in AD were enriched for transcription factor DNA-binding motifs. *De novo* motifs identified by HOMER were matched to transcription factors using Tomtom and promoter histone acetylation (**Table S9**). The top motif enriched at microglia hyperacetylated regions was for the MITF/TFE family and other bHLH-ZIP, E-box-binding transcription factors (**Figure 5B**). Interestingly, several *de novo* motifs were linked to transcription factors that were directly hyperacetylated in microglia, including *MITF, NFE2, BACH1, KLF6*, *SREBF1* and AP-1 family members, indicating that the expression of these factors is elevated in AD and that they regulate a programme of AD microglial genes (**Figure 5C**). Lysosomal-associated MITF is a regulator of the DAM substate, AP-1 activation (FOS, FOSL2, BATF) is enriched in activated microglia^8^, and KLF6 is a pro-inflammatory factor in macrophages^41^. SREBF1 is a master regulator of fatty acid synthesis genes, in line with lipid metabolism dysregulation in DAM^42^. NFE2 and BACH1 bind antioxidant response motifs as part of the CNC-bZIP family, indicating activation of an oxidative stress response. Conversely, microglia hypoacetylated regions were proximal to the TEF (thyrotroph embryonic factor) circadian factor, which has an altered cycling amplitude in human AD microglia^43^ (**Figure 5C and S8A**).

*De novo* motifs were identified for hyperacetylated regions in oligodendrocytes for the FOX, NF-κB, E2F/DP, HIF, and NFAT families, which have been linked to oxidative stress and aging, neuroinflammation, cell-cycle control of OPC differentiation^44^, hypoxia signalling, and calcineurin/NFAT-mediated myelination^45^, respectively (**Figure 5D**). FOXO3-associated enhancers were directly elevated in AD (**Figure 5E**). FOXO3 has been associated with human longevity and AD risk through microglia and oligodendrocyte gene networks^46^. Regulatory regions proximal to KLF13 were altered in AD (**Figure 5E**), which has been linked to oligodendrocyte differentiation and myelin gene expression^47^. Whereas oligodendrocyte hypoacetylated regions were enriched for zinc finger transcription factor motifs (**Figure S8B**). Lastly, neuronal hyperacetylated regions were enriched for motifs of the PAR-bZIP circadian family (DBP, HLF, TEF) and members of the Regulatory Factor X (RFX) family, RFX1 and RFX5 (**Figure S8C**). Neuronal hypoacetylation was linked to the MEF2 family (**Figure S8D-E**), consistent with prior evidence linking reduced MEF2 activity to cognitive decline^48^.

### MITF is a key transcriptional regulator in AD microglia

The most enriched motif at microglia hyperacetylated regions in AD was for the MITF/TFE transcription factors, which are associated with lysosomal/autophagy-related pathways. In addition, *MITF* was directly hyperacetylated (**Figure 6A-B**), and the expression of *MITF* has been reported to be elevated in human and mouse AD models^49-51^. However, the genome-wide binding and target genes for MITF in human brain resident microglia have not been reported. MITF binding was determined using cleavage under targets and tagmentation (CUT&Tag) in microglia nuclei from human resected cortical tissue (**Figure S9A-B, Table S10**). The majority of the MITF binding regions overlapped with microglia H3K27ac gene regulatory regions (93%) (**Figure 6C**). Genes proximal to MITF binding sites were associated with the endolysosomal compartment, catabolic processes and vacuole organisation (**Figure 6D**). *De novo* HOMER motif analysis identified MITF/TFE and BHE41 as the top three DNA-binding motifs, collectively indicating that CUT&Tag enriched for microglial MITF binding sites (**Figure 6E**). Additional motifs that were identified at MITF binding regions included SPI1/SPIB, IRF8, C/EBP and AP-1 transcription factor families that may represent collaborative binding partners (**Figure 6E**). PU.1 (*SPI1*) is a known binding partner of MITF in osteoclasts^52^, and MITF co-occupies the same genomic regions as AP-1 transcription factors^53^. MITF binding signal overlapped active gene regulatory elements (**Figure 6C and 6F**) and was more enriched at microglia regions hyperacetylated in AD compared to total microglia H3K27ac (Wilcoxon rank-sum test, p=4e-8) (**Figure 6F**). Lastly, genes proximal to hyperacetylated regions in microglia that intersect with MITF binding sites were enriched for microglia DAM signature genes^25^ (**Figure 6G**). This suggests that MITF may drive the transition to a disease-associated state and aligns with evidence of MITF driving a disease-associated transcriptional signature^54,55^.

**Figure 6.**
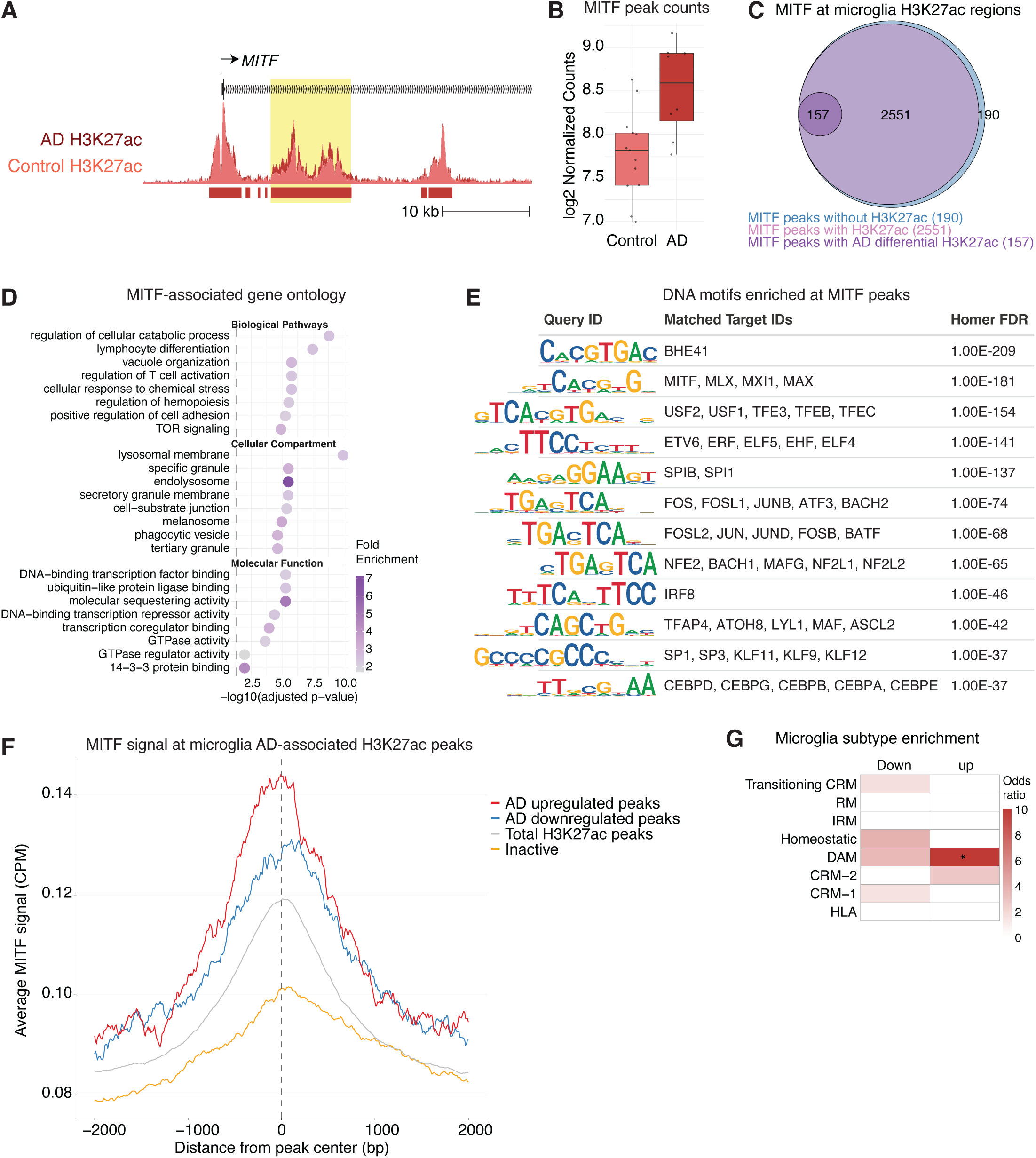
MITF is a key transcriptional regulator in AD microglia. **(A)** UCSC browser visualisation of microglia H3K27ac signal at the *MITF* locus in control (pale red) and AD (dark red). Yellow highlight, differential H3K27ac region. **(B)** Read counts per million for the microglia *MITF* peak with differential H3K27ac in AD. **(C)** Venn diagram of microglia MITF CUT&Tag peaks that overlap with microglia H3K27ac regions. **(D)** Gene ontology enrichment analysis of genes annotated to microglia human MITF CUT&Tag peaks using enrichGO. **(E)** *De novo* motif analysis of human microglia MITF binding regions. *De novo* motifs were matched to transcription factors using Tomtom and filtered on transcription factors with active promoters in microglia. **(F)** Aggregate plot of mean MITF CUT&Tag signal at H3K27ac peaks hyperacetylated (red) and hypoacetylated in AD (blue), total H3K27ac peaks (grey) and inactive promoters (yellow) in microglia. **(G)** Fisher’s exact test for enrichment of annotated microglial differential H3K27ac peaks that overlap with MITF binding sites for signature genes of microglial transcriptional substates defined using a human-mouse amyloid model by scRNA-seq^25^.

## Discussion

Histone acetylation profiles in microglia, oligodendrocytes and neurons revealed coordinated shifts in regulatory states in late-stage AD. Sub-cell-state analyses indicated a shift of microglia from homeostatic to an activated, disease-associated state, as well as changes in oligodendrocyte precursor cells and mature oligodendrocytes, and vulnerability of excitatory and inhibitory neuronal populations. Integration of genetic risk with allele-specific H3K27ac analyses highlighted AD risk variants that potentially perturb regulatory regions in microglia and oligodendrocytes. Finally, hyperacetylation of the lysosomal regulator *MITF*, together with enrichment of the MITF/TFE recognition motif and DNA binding of MITF in microglial hyperacetylated regions, implicated MITF as a candidate transcriptional driver of a DAM cell state.

Microglial hypoacetylation at homeostatic and hyperacetylation at inflammatory, lipid processing and phagocytic signature genes aligns with observations that at later disease stages, specific microglial states are associated with amyloid plaques^56-58^. Human gene expression studies have identified corresponding DAM and amyloid-responsive microglia states that are plaque-proximal and modulated by AD risk alleles^51,59,60^. Co-regulated gene regulatory regions that positively correlated with AD included signature genes of monocyte-derived macrophages, which is consistent with recent findings of brain infiltration of monocyte-derived macrophages with age^61,62^ and suggests this may be elevated further in disease. Neuronal hypoacetylation was enriched for signature genes of vulnerable SST⁺ and RELN⁺/LAMP5 interneurons and RELN⁺ EC layer II/III excitatory neuronal populations that are depleted in AD yet relatively preserved in resilient individuals^1,3,63^. Oligodendrocytes showed the most extensive H3K27ac dysregulation in AD, affecting immune and neuron-oligodendrocyte interaction signalling pathways. These changes are consistent with widespread transcriptomic and epigenomic remodelling of oligodendrocytes reported in late-stage AD^12,64^.

H3K27ac ASVs in oligodendrocytes and microglia coincided with AD risk variants at the *BIN1* and *PICALM* loci in microglia and oligodendrocytes, as well as at the *CLU* locus in oligodendrocytes and *HLA-DRB1*, *APOE*, *CASS4* and *MS4A4* loci in microglia. *BIN1* expression is increased in the AD brain^65^, and microglial *BIN1* has been implicated in the spreading of tau^66^. Fine-mapping and promoter-anchored chromatin contacts linked a high-confidence AD variant (rs6733839) in a microglia-specific enhancer to the *BIN1* promoter^19,67^. The *BIN1* lead variant has been identified as a macrophage chromatin accessibility quantitative trait locus^68^, and CRISPR deletion of the enhancer abolishes *BIN1* expression in iPSC-derived microglia but not in neurons or astrocytes^16^. Similarly, a high-confidence AD variant (rs10792832) has been localised to a microglia-specific enhancer at the *PICALM* locus^16^. This variant was associated with reduced PU.1 binding, lower *PICALM* expression, altered cholesterol synthesis and impaired phagocytosis of amyloid beta and myelin^69^. In alignment, *APOE* alleles have been linked to amyloid pathology, cholesterol metabolism^70,71^, and phagocytosis^60^; and *MS4A4A* to negative regulation of TREM2 signalling, lipid metabolism and a chemokine microglial cell state^72,73^. Human Leukocyte Antigen DR (HLA-DR) is a component of Major Histocompatibility Complex Class II (MHCII) presentation and is highly expressed on reactive microglia in AD brains^74^.

The lysosomal regulator MITF was identified as a key transcriptional regulator of AD microglia. The *MITF* gene was directly hyperacetylated, and the MITF/TFE recognition motif and DNA binding of MITF were enriched across microglia hyperacetylated regions. This aligned with the identification of microglial ASVs and AD hyperacetylated regions proximal to endolysosomal and vesicle-associated genes. Lysosomal function in microglia has been implicated in immune activation, and clearance of damaged organelles and toxic protein aggregates^75,76^. Gene expression studies identified *MITF* as a core neurodegeneration microglia hub gene across AD models^49^ and in human AD brains^50^. *MITF* overexpression in iPSC-derived microglia-like cells (iMGL) induced a DAM gene expression signature and enhanced phagocytosis^54^. Furthermore, treatment of iMGL with histone deacetylase inhibitors upregulated *MITF*, induced a DAM-like signature, increased amyloid uptake and reduced secretion of monocyte chemoattractant protein-1 (MCP-1)^77^. Recently, a single-cell gene expression analysis of human myeloid cells implicated MITF as an upstream regulator of an AD-associated microglial subtype with elevated *GPNMB^55^*. Overall, our findings now provide human *in vivo* epigenomic evidence that lysosomal-associated gene regulation is altered in microglia in individuals with AD.

Collectively, these results highlight the importance of glial gene regulatory changes in disease and implicate dysfunction of lysosomal-associated pathways in brain immune cells in AD, providing a framework for target prioritisation. Future studies with increased sample size will provide a deeper characterisation of allelic effects on regulatory landscapes for fine mapping of causal variants. Applying single-cell epigenomic analysis will enable higher-resolution analysis of glial states and neuronal subtypes, and analysis of additional cell types such as vascular-associated brain endothelial cells, mural cells and astrocytes, which were recently associated with genetic risk for dementia^17,78,79^.

## Author Contributions

A.N. and C.K.G. conceptualised the project. Resources (tissue) were provided by P.N., R.A.R., and J.B.B. Wet lab methodology was developed by A.N. and J.C.M.S. Investigation (data collection) was performed by A.N., C.K.N., M.P.P. and K.C.Z. Computational data curation was performed by C.G. and P.W., and analysis and visualisation were performed by C.G., P.W., P.U., Y.W., J.L.T., A.A., and R.M.Y., with contributions from I.R.H. and S.J.M. Manuscript writing was by A.N., C.G., P.W., and Y.W., with contributions from all authors. All authors reviewed and edited the manuscript.

## Acknowledgments

We would like to thank Anna Mallach, Sam Boulger, and members of Nathan Skene’s (Imperial College London) and Dervis Salih’s (University College London) groups for advice and discussion. We would like to thank Nik Matthews and his team at Imperial BRC Genomics and the UCSD sequencing facility for their help with sequencing.

## Resource availability

Data is available as fastq files on GEO [GSE341557]. Code is available at https://github.com/Nott-Group/AD_epigenome and https://github.com/Marzi-lab/allelic_H3K27ac. Peak coordinates and interactomes are provided in Supplemental Tables. A UCSC genome browser session (hg38) containing the processed H3K27ac ChIP-seq datasets for each brain cell type is located at: https://nottgroup.com/resources.html.

## Supplementary Figure Legends

**Figure S1.**
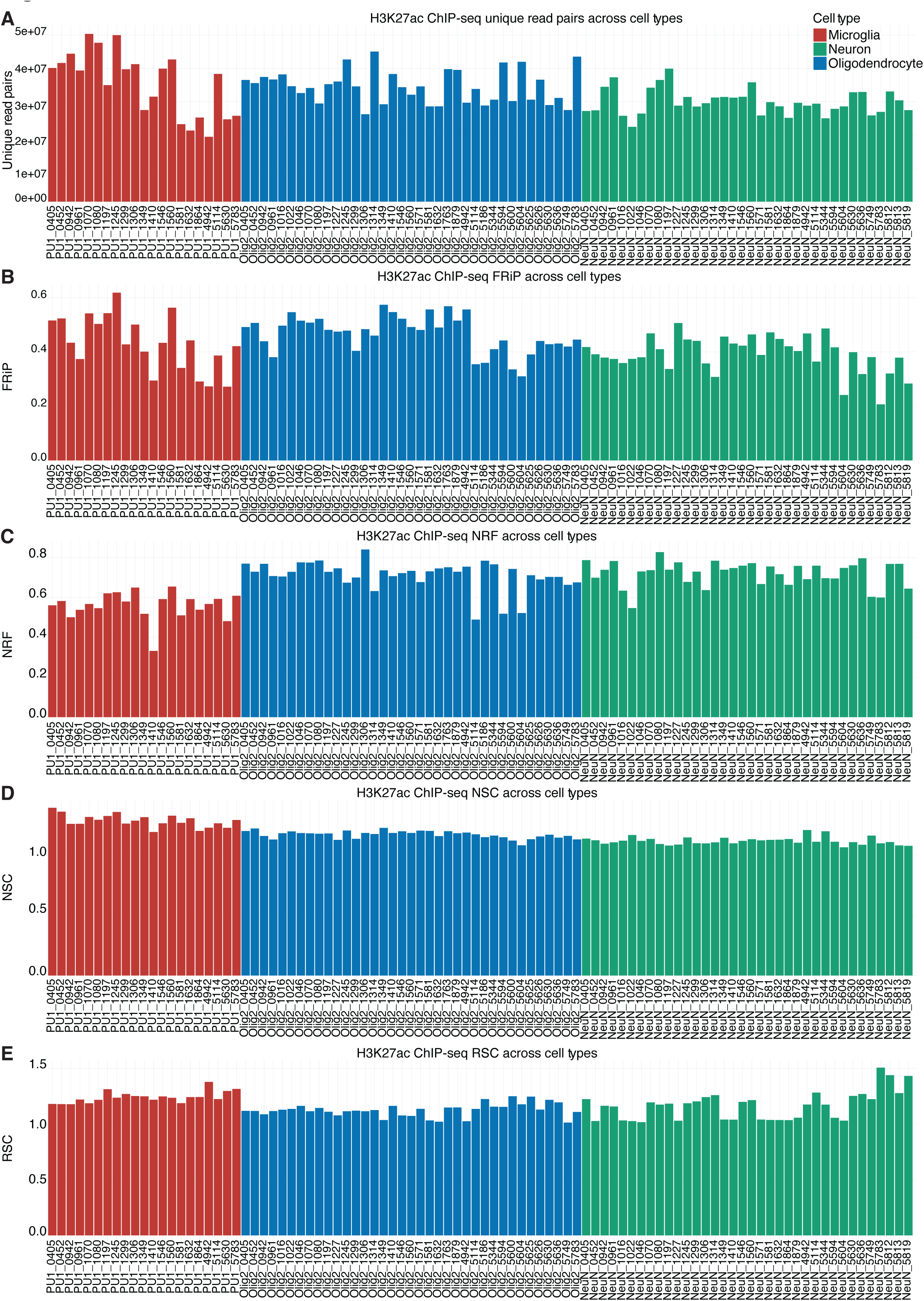
Quality control metrics for H3K27ac ChIP-seq. **(A)** Number of unique reads for microglia, oligodendrocytes and neuron samples. **(B)** FRiP scores for microglia, oligodendrocytes and neuron samples. **(C)** Non-redundant fraction (NRF) as a measure of duplicates for microglia, oligodendrocytes and neuron samples. **(D)** Normalised strand coefficients (NSC) for microglia, oligodendrocytes and neuron samples. **(E)** Relative strand correlation (RSC) for microglia, oligodendrocytes and neuron samples.

**Figure S2.**
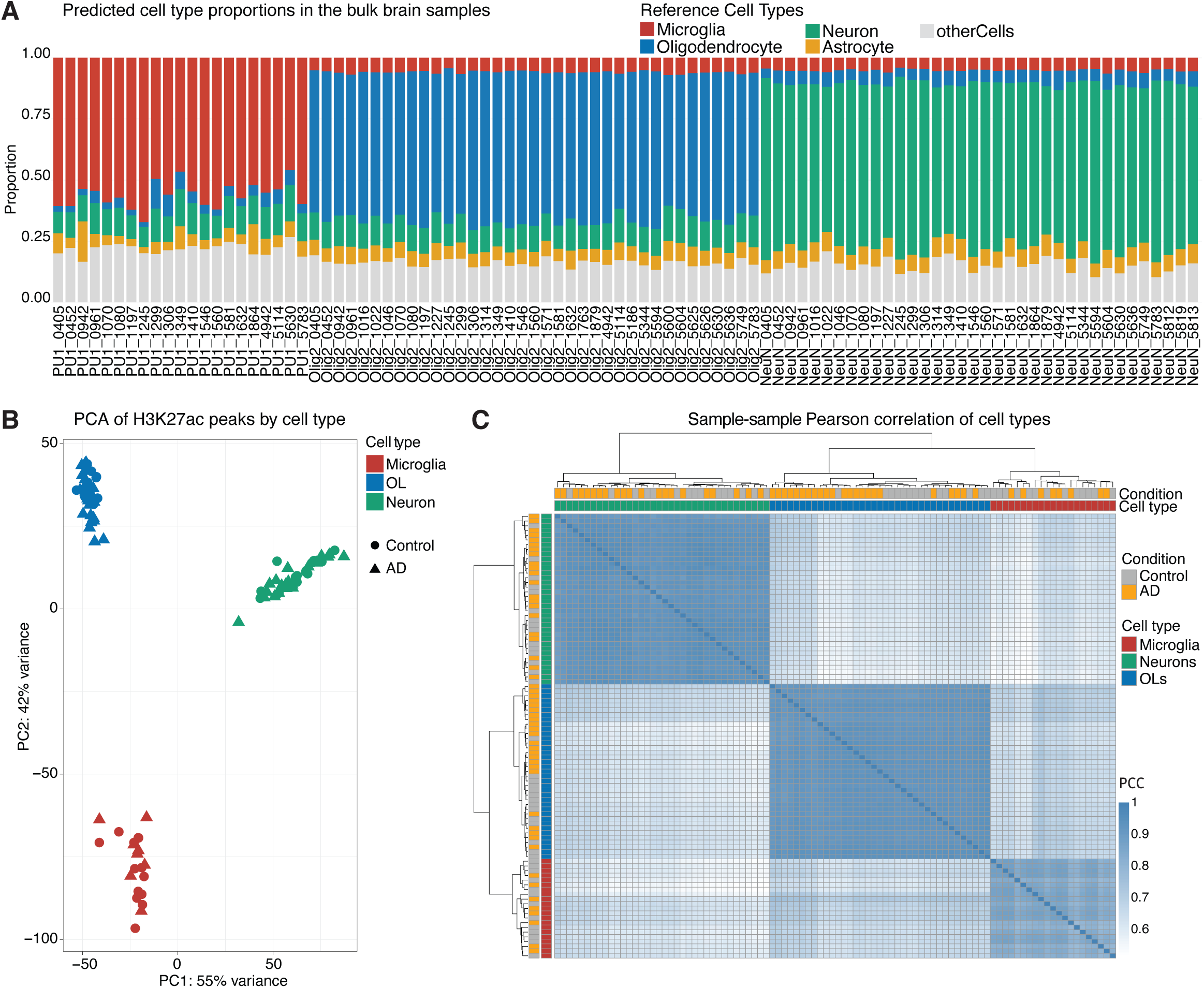
Cell type enrichment of H3K27ac ChIP-seq samples. **(A)** Cell type proportion estimates using CHAS for H3K27ac ChIP-seq samples of nuclei enriched for microglia, oligodendrocytes and neurons. **(B)** Principal component analysis on variance-stabilised counts using DESeq2. **(C)** Pearson’s correlation heatmap of H3K27ac ChIP-seq samples for microglia, oligodendrocytes and neurons.

**Figure S3.**
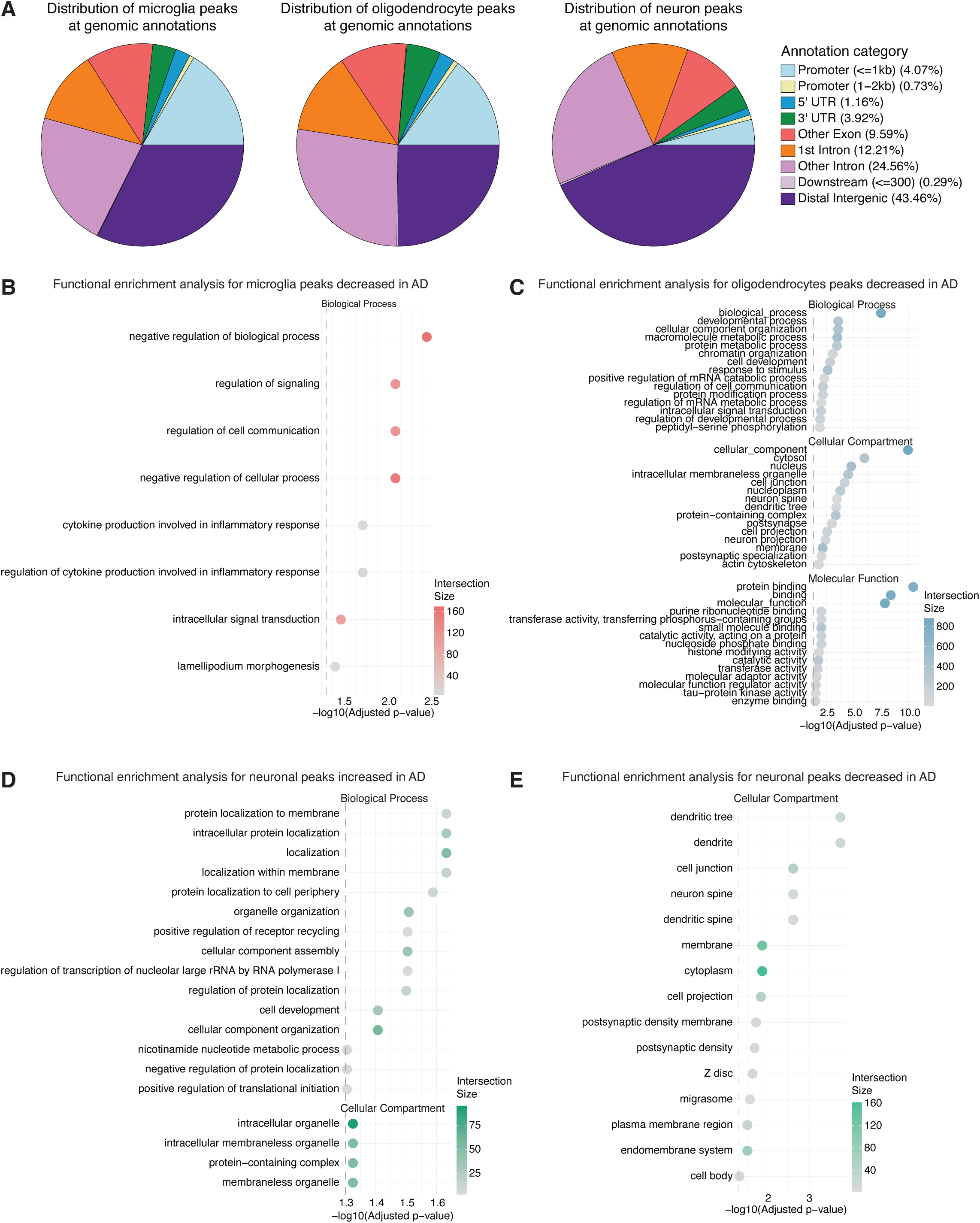
Cell type differential acetylation in AD. **(A)** Pie graph of differential peak annotations (promoter, exon, intron, intergenic, UTRs, CpG islands) for microglia, oligodendrocytes and neurons. **(B)** Functional enrichment analysis of annotated hypoacetylated regions in microglia using gprofiler. **(C)** Functional enrichment analysis of annotated hypoacetylated regions in oligodendrocytes using gprofiler. **(D)** Functional enrichment analysis of annotated hyperacetylated regions in neurons using gprofiler. **(E)** Functional enrichment analysis of annotated hypoacetylated regions in neurons using gprofiler.

**Figure S4.**
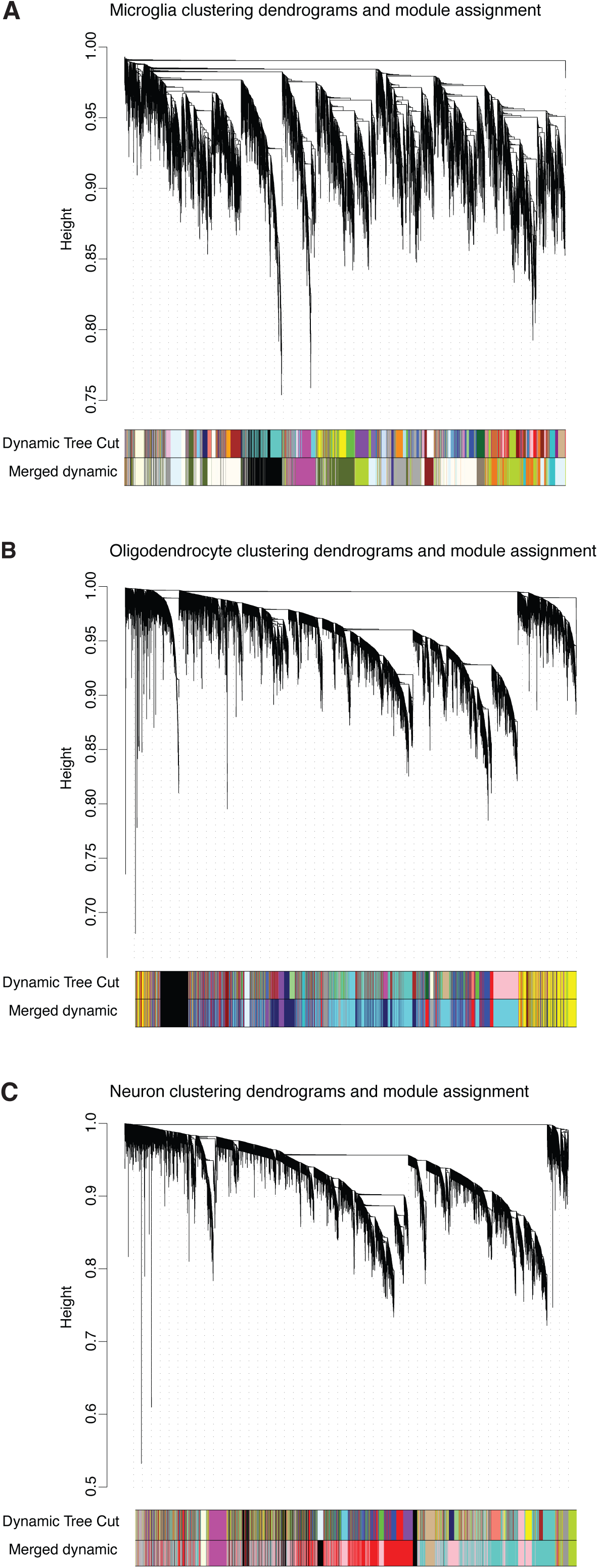
Weighted gene co-expression network analysis. **(A)** Hierarchical clustering dendrogram of microglia H3K27ac promoter peaks based on topological overlap. **(B)** Hierarchical clustering dendrogram of oligodendrocyte H3K27ac promoter peaks based on topological overlap. **(C)** Hierarchical clustering dendrogram of neuron H3K27ac promoter peaks based on topological overlap. Genes are grouped by similarity, with more closely related genes joined lower in the tree. Colour bars below the dendrogram show module assignments from the initial Dynamic Tree Cut (top) and after merging similar modules based on eigengene correlation (Merged dynamic, bottom).

**Figure S5.**
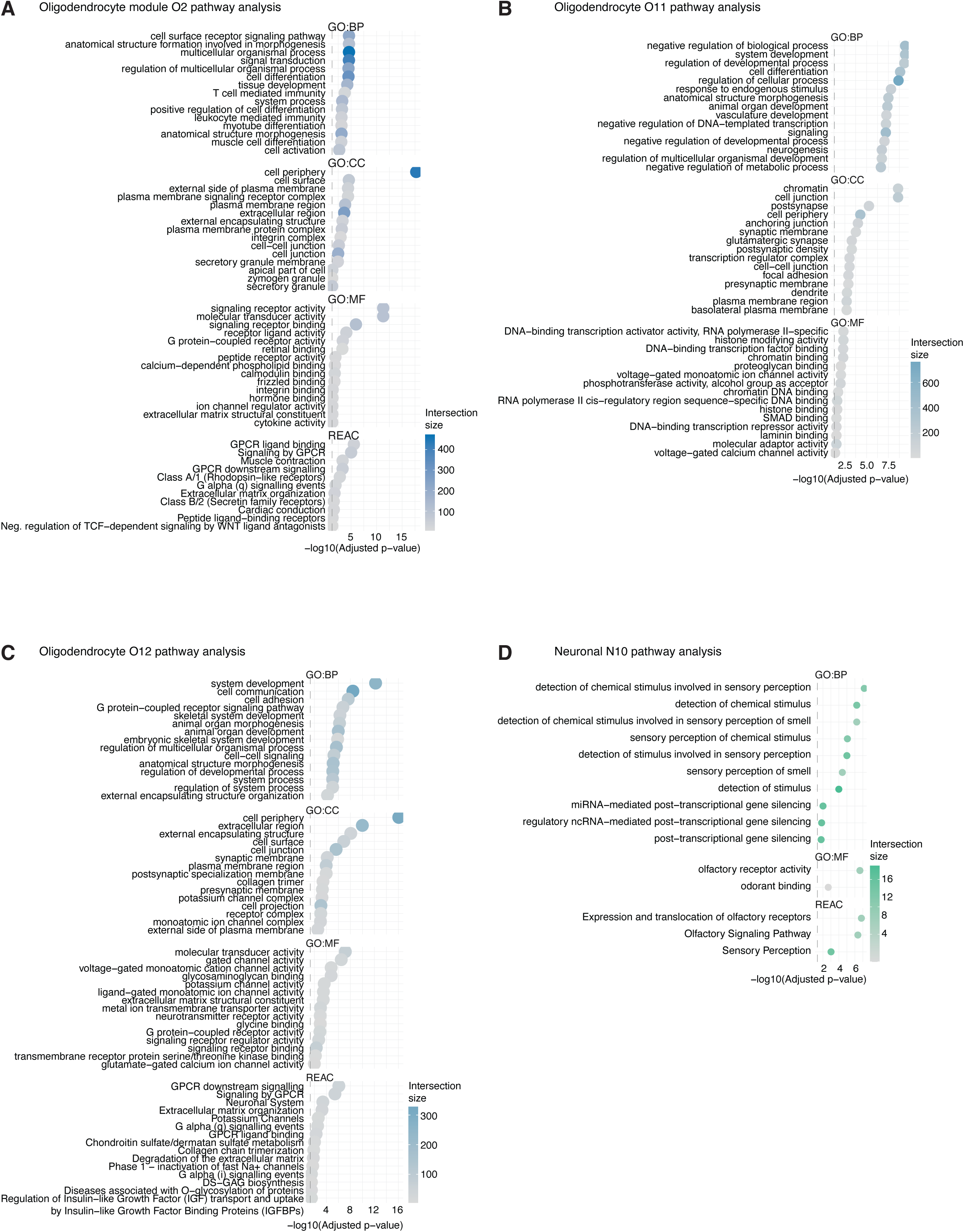
Pathway analysis of WGCNA modules. **(A)** Functional enrichment analysis of oligodendrocyte module O2 genes using gprofiler. **(B)** Functional enrichment analysis of oligodendrocyte module O11 genes using gprofiler. **(C)** Functional enrichment analysis of oligodendrocyte module O12 genes using gprofiler. **(D)** Functional enrichment analysis of neuron module N10 genes using gprofiler.

**Figure S6.**
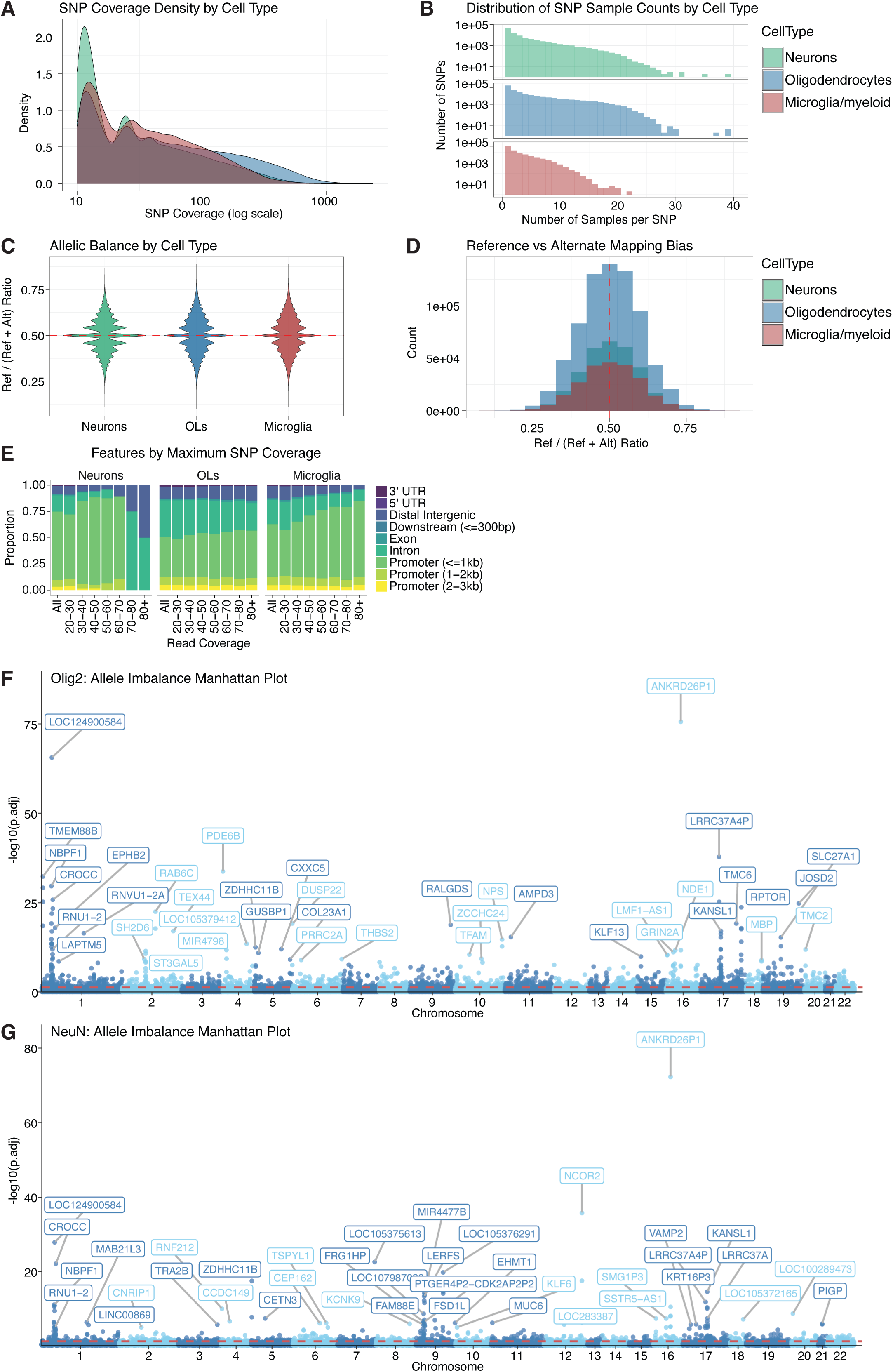
Allele-specific variant analysis across brain cell types. **(A)** SNP coverage density for microglia, oligodendrocytes and neurons. **(B)** Distribution of SNP sample counts by cell types. **(C)** Allelic balance by cell types as ratio of reference/(reference + alternative). **(D)** Mapping bias of reference vs alternative allele as counts. **(E)** Proportion of features (promoter, exon, intron, intergenic, UTR) using ChIPseeker by maximum SNP coverage for microglia, oligodendrocytes and neurons. **(F)** Manhattan plot of oligodendrocyte allele-specific variants with the top 40 annotated to the proximal gene. **(G)** Manhattan plot of neuron allele-specific variants with the top 40 annotated to the proximal gene.

**Figure S7.**
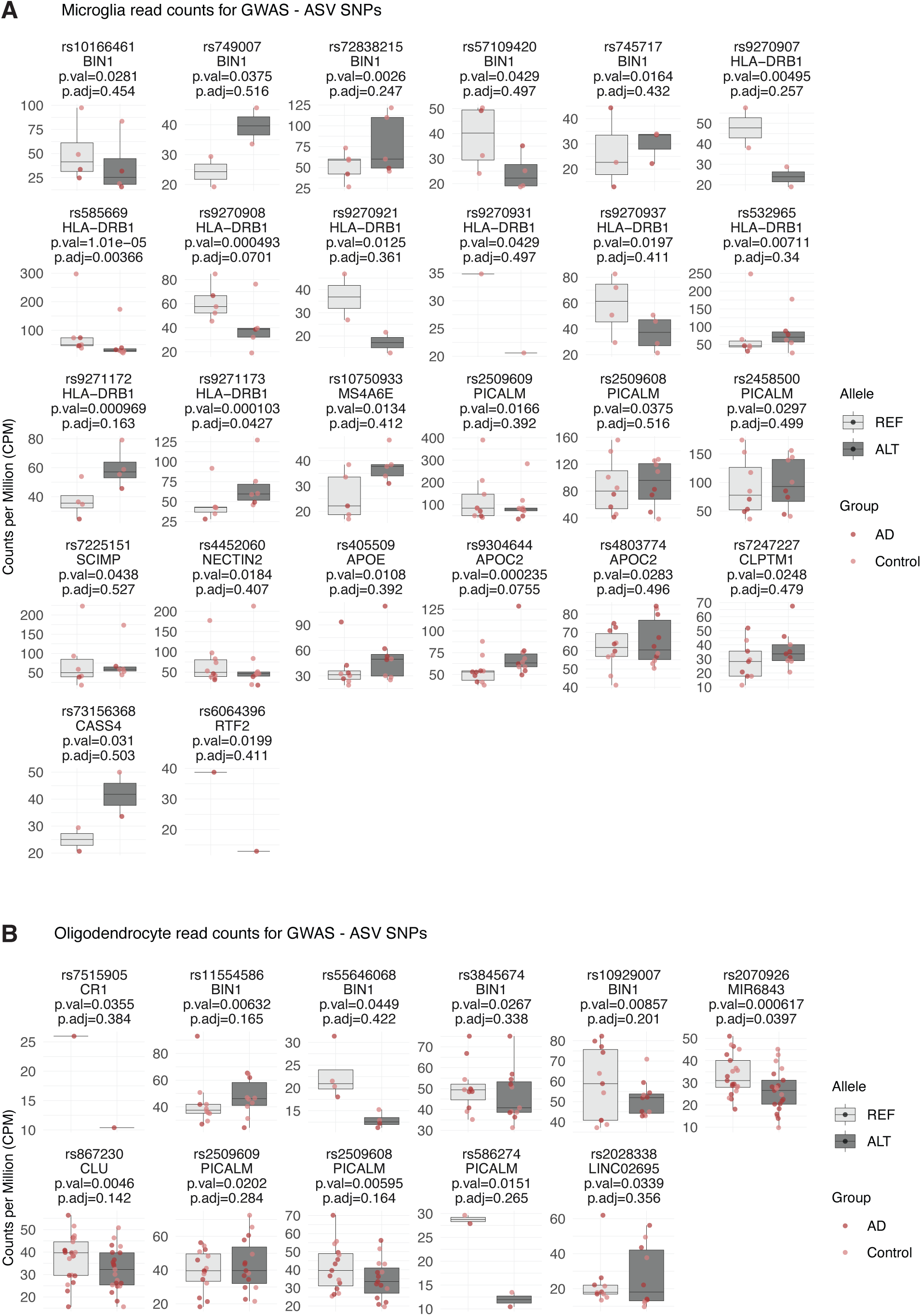
Read counts for allele-specific variants at AD GWAS risk loci. **(A)** Microglia read counts for reference and alternative alleles of two significant allele-specific variants (rs9271173 and rs585669) and nominally significant allele-specific variants at AD GWAS loci. **(B)** Oligodendrocyte read counts for reference and alternative alleles of a significant allele-specific variant (rs2070926) and nominally significant allele-specific variants at AD GWAS loci.

**Figure S8.**
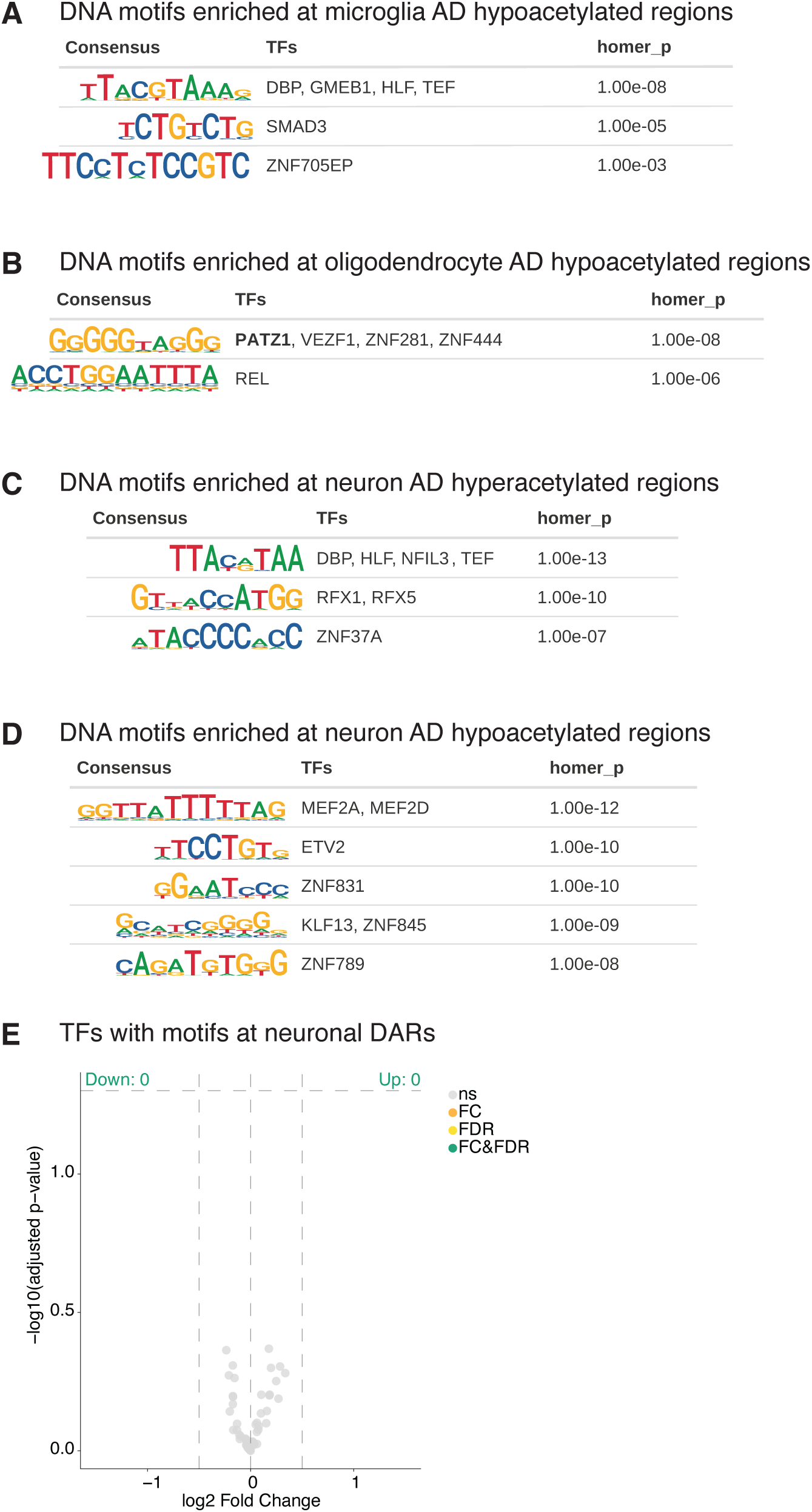
Transcription factor binding site motif matching reveals putative drivers of AD epigenomic dysregulation. **(A)** *De novo* motif analysis of microglia H3K27ac regions hypoacetylated in AD. *De novo* motifs were matched to transcription factors using Tomtom and filtered on transcription factors with active promoters in microglia. **(B)** *De novo* motif analysis of oligodendrocyte H3K27ac regions hypoacetylated in AD. *De novo* motifs were matched to transcription factors using Tomtom and filtered on transcription factors with active promoters in oligodendrocytes. Motif in bold is matched to a transcription factor with proximal differentially acetylated regions in AD. **(C)** *De novo* motif analysis of neuron H3K27ac regions hyperacetylated in AD. *De novo* motifs were matched to transcription factors using Tomtom and filtered on transcription factors with active promoters in neurons. **(D)** *De novo* motif analysis of neuron H3K27ac regions hypoacetylated in AD. *De novo* motifs were matched to transcription factors using Tomtom and filtered on transcription factors with active promoters in neurons. **(E)** Volcano plot of differential H3K27ac regions annotated to transcription factors with motifs enriched at neuronal H3K27ac regions hyperacetylated and hypoacetylated in AD.

**Figure S9.**
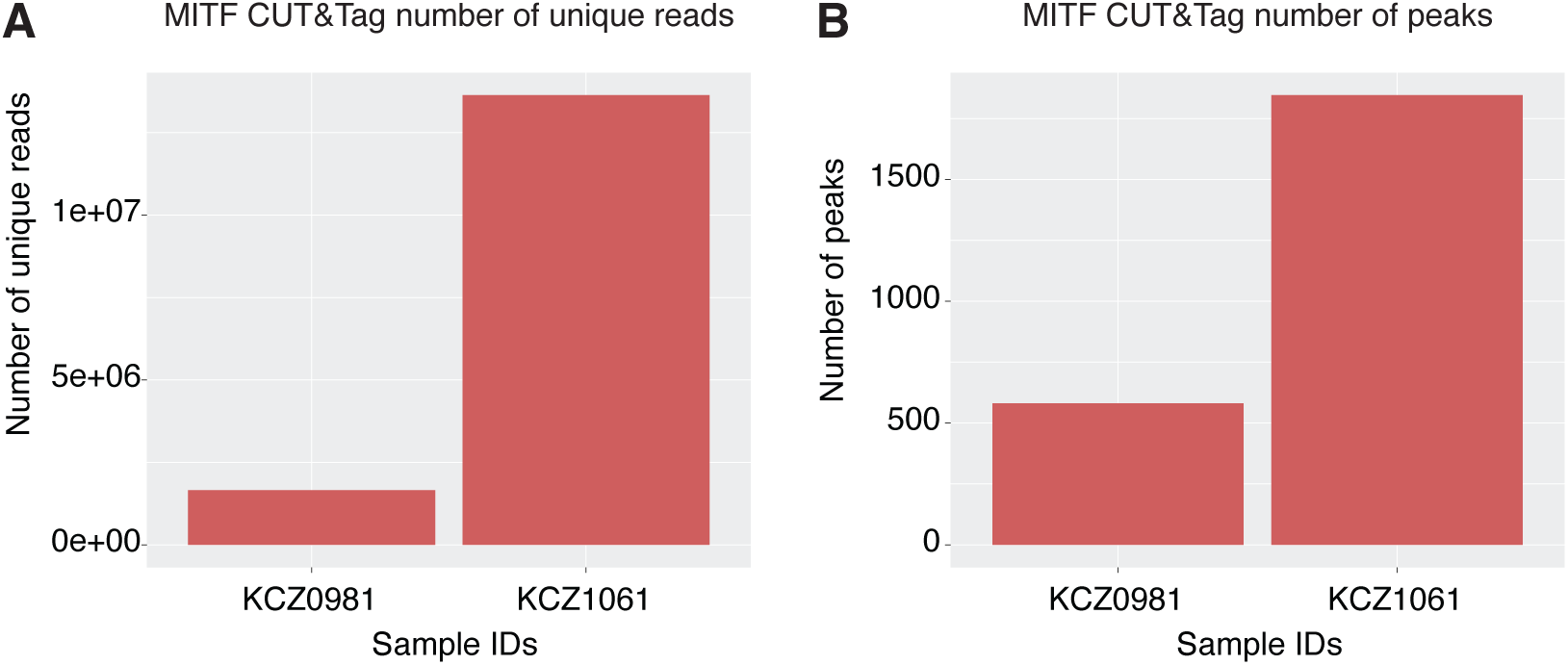
MITF CUT&Tag library metrics. **(A)** Number of unique reads per sample for MITF binding sites identified in microglia through transcription factor CUT&Tag. **(B)** Number of peaks called per sample for MITF binding sites identified in microglia through transcription factor CUT&Tag.

## Supplemental Tables

**Table S1.** Sample metadata

**Table S2.** H3K27ac consensus peaks

**Table S3.** H3K27ac peak featureCounts

**Table S4.** Differential H3K27ac DESeq2 output

**Table S5.** Differential peaks pathway analysis

**Table S6.** WGCNA module membership

**Table S7.** WGCNA module pathway analysis

**Table S8.** Allele-specific variant analysis

**Table S9.** Transcription factor matches

**Table S10.** MITF consensus peaks

## Methods

### Sample cohort

Pre-frontal cortex was sourced as fresh frozen tissue blocks from the University of California, San Diego (UCSD) Shiley-Marcos Alzheimer’s Disease Research Center (Broadmann area (BA)9, 46 and 10), the University of Kentucky AD Research Center (UK-ADRC) autopsy cohort biobank (BA9), and the Banner Sun Health Research Institute Brain and Body Donation program (BA9) (**Table S1**). Donors with late-stage AD were selected with Braak stage V-VI, and non-dementia controls with Braak stage 0-II (**Table S1**). Donors were aged between 59 and 94, matched for sex (**Table S1**). All tissue samples were stored at -80 °C, and nuclei were FANS-isolated at UCSD. H3K27ac ChIP-seq was performed on nuclei samples at UCSD and Imperial College London (**Table S1**).

The UK-ADRC is a community-based cohort recruiting from the Lexington, Kentucky region^81^. Protocols were approved by the University of Kentucky Institutional Review Board, and all participants provided written informed consent.

### Cell-type nuclei isolation

Nuclei were enriched according to the cell type of origin by FANS as previously described with minor modifications^23^. Approximately 250 mg tissue was homogenised in 0.3 ml 1% formaldehyde in phosphate-buffered saline (PBS) using a pellet pestle and motor (Kimble, #KT749521-0590, #KT749540-0000). Homogenates were adjusted to a volume of 10 ml 1% formaldehyde in PBS and rocked at room temperature for 10 mins. Homogenates were quenched with 0.5 ml 2.5 M glycine and rocked at room temperature for 5 mins. Homogenates were pelleted at 1,100xg for 5 mins at 4^°^C and washed two times with 10 ml NF1 buffer (10 mM Tris-HCl, pH 8.0, 1 mM EDTA, 5 mM MgCl2, 0.1 M Sucrose, 0.5% Triton X-100). Samples were resuspended in 5 ml NF1 after final wash and incubated on ice for 5 mins. Samples were dounce homogenised using 20 strokes of a loose pestle and 5 strokes of a tight pestle (Wheaton, ThermoFisher #06435A). Homogenates were passed through a 70 μm strainer and washed through with 15 ml NF1 (20 ml total). Samples were underlaid with 5 ml 1.4 M sucrose cushion (40% sucrose, 10 mM Tris-HCl, pH 8.0, 3 mM MgCl2, 1 mM DTT) and centrifuged at 3,900xg for 30 mins at 4°C. Pelleted nuclei were washed once with 10 ml NF1 and once with staining buffer (1% BSA, 1 mM EDTA in PBS) at 1,600xg for 5 mins at 4°C. Pelleted nuclei were resuspended in 0.6 ml staining buffer, and 1% was retained for a non-stained control. Nuclei were stained with anti-NeuN-AF488 (1:2,500, Sigma-Aldrich #MAB377X), anti-PU.1-PE (1:100, Cell Signaling Technology #81886S), and anti-OLIG2-AF647 (1:1,000, Abcam #ab225100) at 4°C overnight. The next day, nuclei were washed with 4 ml staining buffer at 1,600xg for 5 mins at 4°C and resuspended in 0.5 ml staining buffer. Nuclei were passed through a 30 μm strainer and adjusted to 4 ml with staining buffer. Nuclei were counterstained using 0.5 µg ml^−1^ DAPI and enriched according to cell type using a Beckman Coulter MoFlo® Astrios™ EQ cell sorter. Cell type-enriched nuclei were pelleted at 1,600xg for 15 mins in staining buffer and stored at -80°C until processing for ChIP-seq.

### Chromatin immunoprecipitation followed by sequencing

ChIP-seq was performed as described with minor modifications^16,82^. Nuclei pellets were thawed on ice, resuspended in 130 μl LB3 (10 mM Tris-HCl pH 7.5, 100mM NaCl, 1 mM EDTA (pH 8.0), 0.5 mM EGTA, 0.1% Na-deoxycholate, 0.5% N-laurosylsarcosine, 1X protease inhibitor cocktail (PIC)) and transferred to Covaris microTubes (Covaris #520045). Pellets were sonicated using a Covaris E220 (duration, 10 x 60 secs; duty: 5.0; PIP, 140; cycles: 200; amp/vel/dwell: 0.0). Sonicated samples were transferred to 1.5 ml tubes and volumes adjusted to 250 μl with LB3 and 25 μl 10% Triton 100-x. Samples were centrifuged at maximum speed for 10 mins at 4°C and transferred to new 1.5 ml tubes. 1% of each sample was aliquoted for input controls and stored at -20°C. Microglia samples with ∼100,000 nuclei were supplemented with the carriers: 5 μg H2B (NEB cat # M2505S) and 0.25 μg spleen mRNA (Zyagen cat # MR-701-MR). 25 μl Protein G DynaBeads (Life Technologies, #10004D) and 2 μl anti-H3K27ac polyclonal antibody (ActiveMotif, #39135) were added to each sample and rotated at 4°C overnight. The next day, samples were placed on a magnet and supernatant removed. DynaBeads were resuspended with ice-cold 150 μl WB1 (20 mM Tris-HCl pH 7.4, 150 mM NaCl, 2 mM EDTA (pH 8.0), 0.1% SDS, 1% Triton X-100, 1X PIC) and transferred to PCR tubes. Samples were passed across a magnet 10 times, the wash buffer was removed, resuspended with 150 μl WB1 and transferred to new PCR tubes. The WB1 wash was repeated for a third time. DynaBeads were washed three times with 150 μl WB3 (10 mM Tris-HCl pH 7.5, 250 mM LiCl, 1% Triton X-100, 1 mM EDTA (pH 8.0), 0.7% Na-deoxycholate) and three times with 150 μl TET (10 mM Tris pH 7.5, 1 mM EDTA, 0.2% Tween 20). DynaBeads were transferred to new PCR tubes on the third TET wash and washed once with 150 μl TE-NaCl (10 mM Tris pH 7.5, 1 mM EDTA, 50 mM NaCl). DynaBeads were resuspended in 25 μl TT (10 mM Tris pH 8.0, 0.05% Tween20). Input controls from day one were thawed and 22.5 μl TT was added.

DNA libraries were prepared using the NEBNextRTM Ultra II DNA Library Prep Kit (New England Biolabs, #E7645L) as follows: 3.5 μl Green NEBNext Ultra II End Prep Reaction Buffer and 1.5 μl Green NEBNext Ultra II End Prep Enzyme Mix were added to 25 μl samples, briefly vortexed and incubated at 20°C for 30 mins and 65°C for 30 mins. 15 μl Red NEBNext Ultra II Ligation Master Mix, 0.5 μl Red NEBNext Ultra II Ligation Enhancer and 1 μl diluted NEXTFLEX Unique Dual Index Barcodes (diluted 1:50, BIOO Scientific, #NOVA-514150) were added and incubated at 20°C for 20 mins. 16 μl H2O, 4 μl 10% SDS, 3 μl 0.5 M EDTA, 4 μl 0.2 M EGTA, 1 μl 20 mg ml^-1^ proteinase K (NEB # P8107S), 1 μl 10 mg ml^-1^ RNase A (Sigma # R5000), and 4.5 μl 5 M NaCl were added to each sample. Samples were briefly vortexed and incubated at 55°C for 1 hour and 65°C overnight. Next day, the samples were placed on a magnet, samples were transferred to new tubes, and DynaBeads were discarded. SpeedBeads (2 μl) (Life Sciences #GE 65152105050250) were washed once in 200 μl EDTA and once in 200 μl TE. Washed SpeedBeads were resuspended in 124 μl 20% PEG/1.5M NaCl and added to each sample and incubated at room temperature for 15 mins. Beads were washed two times with 80% EtOH, air-dried and eluted with 13 μl 0.5x TT buffer. Samples were placed on a magnet, eluted and transferred to new tubes (beads were discarded). The 13 μl libraries were PCR amplified by adding 12.5 μl NEBNext High-Fidelity 2x PCR MasterMix, 0.125 μl 100 μM Solexa IGA primer, 0.125 μl 100 μM Solexa IGB primer and incubated 98°C for 30 secs, and 14 cycles of 98°C for 10 secs, 60°C for 25 secs, and 72°C for 30 secs, followed by 72°C for 5 mins. Samples were size-selected at 200-500 bp using 10% Tris-borate-EDTA (TBE) gels (Life Technologies #EC62752BOX). Neuron and oligodendrocyte samples were single-end sequenced for 50 cycles and microglia samples were paired-end sequenced for 50 cycles on a HiSeq4000 (Illumina, San Diego, CA) at UCSD IGM Genomics Center and a HiSeq2000 (Illumina, San Diego, CA) at NIHR Imperial BRC Genomics Facility.

### Preprocessing of ChIP-seq fastq files

Demultiplexed fastq files were pre-processed using the nf-core Nextflow pipeline (v2.2.0)^83^. Nf-core preprocessing steps included: generation of standard sequencing data quality control (QC) metrics with FastQC; trimming of adapters using TrimGalore; alignment of reads to GRCh38 reference genome using Bowtie2; marking duplicate reads using Picard; merging alignments of multiple libraries from the same sample using Picard; filtering of reads that are unmapped, multimapped, duplicates, mapping to blacklisted regions, containing more than 4 mismatches using Samtools^84^, Bedtools^85^, and Bamtools^86^. Alignment level QC and generation of library complexity scores were generated using Picard and Preseq, respectively. Strand cross-correlation was generated using Phantompeakqualtools.

### Quality control of processed samples

Sample quality was guided by ENCODE guidelines for histone ChIP-seq^87^. QC metrics were as follows: normalised strand coefficient (NSC) > 1.05, relative strand coefficient (RSC) > 0.8, fraction of reads in peaks (FRiP) score > 0.2. Only samples with a minimum of 19 million unique reads were kept. In addition, we applied a H3K27ac deconvolution tool called cell type specific histone acetylation score (CHAS)^88^, to estimate cell type proportions. CHAS was used as a metric of nuclei enrichment purity. CHAS determines the similarity of samples to prior published cell type-specific H3K27ac ChIP-seq data^16^. Samples needed a CHAS score of 0.4 for the cell type which they are supposed to be enriched for to be retained, i.e. PU.1 samples all had a CHAS Microglia score over 0.4.

### Generation of cell-type peak sets

BAM files for samples that passed QC were used to generate peaks within each cell type. BAM files were first divided into sets derived from AD and control donors, and merged BAMs for control and AD cases were produced separately per cell type. Narrow peaks were called on merged bam files using MACS2 callpeak with -g 2701495711 -s 50 --keep-dup all and -f BAM for oligodendrocyte and neuron samples, and -f BAMPE for microglia samples. This generated 6 peak sets in total, across 3 cell types and 2 conditions. AD and control peaks were then merged for each cell type to create 3 cell type consensus peak sets (microglia, oligodendrocyte and neurons) using GenomicRanges R packages^89^ using reduce() and nearby peaks were merged (100bp or closer). Peaks in the final cell type consensus sets were retained if they were called as peaks in 1 or more of the individual samples of the same cell type (peaks from individual samples were called using MACS2 as described above).

### Validation of cell type enrichment of nuclei

Validation of cell type enrichment was performed using a unified consensus peak set. The unified consensus peak set was generated by merging the cell type consensus peak sets across all three cell types using GenomicRanges^89^ using reduce() and nearby peaks were merged (100bp or closer). Counts per peak per sample for the unified consensus peak set was calculated using featureCounts^90^ for all samples. Principal component analysis was performed on normalized counts using DESeq2^91^. To identify peaks enriched in each cell type, we performed a pairwise differential analysis comparing each cell type to every other cell type. Peaks that were differential in pair-wise comparisons across all cell types were retained as cell type specific peaks. Peaks were annotated to genes with ChIPseeker and peaks within 2000 bp upstream or 500 bp downstream of a transcription start site were considered promoters as assigned by TxDb.Hsapiens.UCSC.hg38.refGene.

Cell-type-specific promoters were assessed for enrichment of cell type gene signatures using Expression Weighted Celltype Enrichment (EWCE)^92^. EWCE was run using bootstrap_enrichment_test() with a single cell gene expression reference generated from adult human prefrontal cortex^80^ and the cell-type-specific promoters annotated to genes. Statistical significance was determined using p-values generated from the bootstrap distribution of 10000 random gene sets and corrected for multiple testing using Benjamini– Hochberg adjustment.

A competitive gene set test accounting for inter-gene correlation, CAMERA (v1.58.0)^93^, was carried out to identify whether microglia, oligodendrocyte and neuronal nuclei were significantly associated with their reciprocal cell types using single cell gene expression data. Marker genes from a single cell gene expression dataset^80^ were used to compare against the promoter annotations for each of the cell types.

Cell-type-specific promoters were assessed for enrichment of cell type signature gene sets with over representation analysis using enricher from the clusterProfiler R package. Gene sets that contain curated markers for cell types identified in single-cell sequencing studies of human tissue from the MSigDb C8: cell type signature gene sets^94^ were used as reference with input gene lists of annotated cell-type-specific promoters.

### Differential peak analysis in AD

Differential H3K27ac peaks between AD cases and ND controls per cell type. A raw read count matrix was generated for each cell type consensus peak set using featureCounts. Differential H3K27ac peaks between AD and ND controls were identified with DESeq2^91^ using the cell type raw read count matrices. Confounding factors included in the modelling were patient age, sex and brain bank. Analysis was implemented with the standard DESeq command, which estimates size factors, dispersion and then fits models on the normalized counts data. Results were then extracted with Benjamini Hochberg adjusted p-values to account for multiple testing. Significantly differential regions were defined as peaks having p.adj<0.05.

### Peak annotation to genes

All peaks were processed into GRanges objects using GenomicRanges^89^ and then assigned to genes with annotatePeak from ChIPseeker^95^ using reference transcriptome TxDb.Hsapiens.UCSC.hg38.refGene and annotation database org.Hs.eg.db.

To identify active genes in each cell type, promoter regions were overlapped with H3K27ac peaks for each cell type. Promoter regions were defined by extending all transcriptional start sites from TxDb.Hsapiens.UCSC.hg38.refGene 2000bp upstream and 500bp downstream. Active gene promoters were identified for each cell type as promoter regions that overlapped with cell-type H3K27ac peaks and were used to define a list of active genes for each cell type. For downstream analyses, only annotations to these active genes were retained.

### Pathway analysis

Annotations of hyperacetylated and hypoacetylated regions were used to perform gene ontology and pathway enrichment analyses with gprofiler2 gost()^96^. For each cell type, analysis was performed separately for the hyperacetylated and hypoacetylated regions. The background for each analysis was all active genes identified in the relevant cell type, as described above in ‘peak annotation to genes’ section.

For visualisation, to reduce redundancy among enriched terms with overlapping gene sets, terms within each ontology were deduplicated based on pairwise Jaccard similarity of their constituent genes (similarity threshold = 0.6). Terms were ranked by significance and processed iteratively, with each term retained only if its maximum Jaccard similarity to any higher-ranked, already-retained term fell below this threshold; terms exceeding the threshold were considered redundant. Full output including all ontology terms can be found in Table S5.

### Cell subtype enrichment analysis

Genes annotated to AD differential H3K27ac peaks for microglia, oligodendrocytes and neurons were assessed for enrichment of cell subtype signature genes using Fisher’s exact test. Signature genes for microglial transcriptional substates were defined using two gene expression datasets^4,25^; 12 substates were defined using human postmortem brain snRNA-seq (including AD and ND controls)^4^ and 8 substates were defined using a human-mouse amyloid model by scRNA-seq^25^. Signature genes for excitatory (Exc) and inhibitory (Inh) neuronal subtypes, oligodendrocyte precursor cells (OPC), and mature oligodendrocytes (Oli) were defined by an snRNA-seq atlas of adult human prefrontal cortex^1^, including AD and ND controls.

Fisher’s exact test (v4.3.1) was used to test if genes annotated to AD differential H3K27ac peaks were overrepresented for cell subtype signature genes compared to a background set of genes. Fisher’s exact test was performed independently for each cell type, and background peak sets were defined as all active genes in the relevant cell type, as described in the ‘peak annotation to genes’ section. Odds ratios, enrichment ratios, expected overlaps, and p-values were calculated for each cell type comparison. Enrichment ratios were visualized using pheatmap (v1.0.12); p<0.05 was annotated on the heatmaps.

### Co-regulation network analysis of H3K27ac peaks

Weighted gene co-expression network analysis (WGCNA) was adapted to identify H3K27ac peaks that are co-regulated, as previously described^97,98^. Modules of correlated H3K27ac peaks for each cell type were generated using WGCNA R package^99^. H3K27ac peaks for microglia, neurons and oligodendrocytes were retained with CPM>1 in at least two samples. Library size differences were normalized using the TMM method (edgeR), and dispersion was estimated (estimateDisp, robust = TRUE) using a design matrix accounting for sex, tissue source, and library size.

The cell type consensus peaks were then annotated relative to transcription start sites using ChIPseeker (annotatePeak, TSS region −2000/+500 bp) against TxDb.Hsapiens.UCSC.hg38.refGene, and peaks overlapping annotated promoters were retained. Promoter peaks were mapped to gene symbols and for genes with multiple promoter peaks, the peak with the highest total signal across samples was retained, resulting in one H3K27ac value per gene. log2-CPM values (cpm, prior.count = 1) for these gene-collapsed promoter peaks were used as the input expression matrix (samples × genes) for network construction.

WGCNA was performed separately for each cell type. A signed weighted adjacency matrix was constructed using a soft-thresholding power (10 for PU.1, 17 for NeuN, and 14 Olig2). Topological overlap (TOM, signed) was calculated from the adjacency matrix, and the corresponding dissimilarity (1 − TOM) was used for average-linkage hierarchical clustering (flashClust). Modules were defined by dynamic tree cutting (cutreeDynamic, deepSplit = 2, minClusterSize = 60, pamRespectsDendro = FALSE) and assigned module colours (labels2colors). Module eigengenes (MEs) were calculated, and modules with eigengene dissimilarity above threshold (MEDissThres = 0.30 for PU.1; 0.2 for Olig2; 0.15 for NeuN) were merged (mergeCloseModules).

Module-trait relationships were assessed by correlating module eigengenes with AD status (coded AD = 1, control = −1) using Pearson correlation, with significance from the Student asymptotic p-value (corPvalueStudent). Modules with p < 0.05 (excluding the unassigned module) were considered significantly associated with AD status. Within-module connectivity (kWithin) was calculated from the TOM (diagonal set to zero).

### Partitioned heritability (stratified LDSC regression)

Stratified LDSC regression was used to quantify the contribution of cell-type regulatory elements to the SNP heritability of AD. All analyses were conducted using the LDSC software package (https://github.com/bulik/ldsc)^100^. Cell type peaks were first lifted over to hg19 using UCSC liftOver then used to create custom genomic annotations. Custom annotation files were generated per cell type using using make_annot.py command. Annotations were generated using the 1000 Genomes Project Phase 3 European-ancestry reference panel. LD scores were computed for each chromosome using the ldsc.py --l2 command, incorporating the custom annotation files alongside the 1000 Genomes Phase 3 European-ancestry genotype data. A 1 cM window was used for LD computation. The --thin-annot option was used to improve computational efficiency by removing SNPs not included in any annotation.

We applied stratified LD score regression using the ldsc.py --h2 command to assess the enrichment of SNP heritability for AD within the cell-type annotations. GWAS summary statistics for AD^15^ were pre-processed and converted to LDSC format using munge_sumstats.py. The model included the baselineLD v2.2 model as a covariate, which consists of annotations designed to control for known confounding factors (e.g., MAF, conservation, histone marks).

Cell type annotations for each cell type were modelled alongside the baseline model using the --ref-ld-chr flag, specifying all annotation sources across chromosomes 1–22. We accounted for overlapping annotations (--overlap-annot) and used the provided allele frequency (--frqfile-chr) and LD weight files (--w-ld-chr) from the 1000 Genomes Phase 3 dataset, excluding the MHC region. Regression coefficients and p-values were computed for each annotation using the --print-coefficients option.

### Allelic Imbalance

Allele-specific variant analyses were performed to quantify genetic imbalance at heterozygous SNPs within cell-type-specific regulatory regions. SNPs were first filtered to extended cell type–specific peaks (±200 bp) using GATK SelectVariants^101^ (v4.3.0.0) and parameter --select-type-to-include SNP. Biallelic heterozygous SNPs at known sites from the 1000 Genomes Project reference panel^102^ were then identified on a per-sample basis with bcftools^103^ view (v1.15.1) with --max-alleles 2, --min-alleles 2, --types snps, --genotype het. These heterozygous SNPs were provided as input to WASP (v0.3.9)^104,105^ to correct for reference mapping bias, followed by duplicate read removal with Picard MarkDuplicates (v2.27.5). Allelic read counts were obtained using GATK ASEReadCounter^101^ (v4.3.0.0). Statistical testing of allele-specific signals was conducted with MixALime^106^ (v2.23.3) under a negative beta-binomial framework, restricting analysis to sites with a minimum of 5 reads per allele. Heterozygous SNPs were intersected with significant AD GWAS SNPs (p<5e-8) from Jansen et al., 2019^15^, and genomic annotation was performed using the ChIPseeker R package (v1.38.0)^95^ with the UCSC hg38 gene transcript database. The workflow is available on GitHub (https://github.com/Marzi-lab/allelic_H3K27ac).

### Transcription factor motif analysis

To identify transcription factors regulating differentially acetylated regions, motif analysis was performed using HOMER (Hypergeometric Optimization of Motif EnRichment)^107^. Hyperacetylated or hypoacetylated regions for each cell type were intersected with chromatin accessibility regions. Cell type chromatin accessibility regions were taken from snMultiome data from human dorsolateral prefrontal cortex (including individuals with no cognitive impairment, mild cognitive impairment and dementia)^78^ using peaks identified as ‘microglia’ for microglia, ‘inhibitory_neuron’ and ‘excitatory_neuron’ for neurons, and ‘oligodendrocyte’ for oligodendrocytes. As background, all ATAC-seq peaks of the respective cell type were used. HOMER parameters were findMotifsGenome.pl input.bed hg38 output_dir -size given -bg background.bed. HOMER *de novo* motifs were mapped to known transcription factor motifs from the HOCOMOCO v14^108^ database using Tomtom in MEME Suite^108,109^, retaining those with significant similarity (q-value ≤ 0.05). Transcription factors were retained that were identified to have active genes in the same cell type, as described in ‘peak annotation to genes’ section. To identify transcription factors with differentially acetylated genes, annotated differential H3K27ac peaks were subset to transcription factors listed in HOCOMOCO v14 matched using the HGNC symbol.

### MITF CUT&Tag

Bulk nano-CUT&Tag for MITF was performed as described for histone modifications^17,79^, with modifications specific to MITF targeting. FANS-sorted PU.1⁺ myeloid nuclei (50,000 per sample; Cell Signaling Technology, Cat. #81886, rabbit) from two paediatric resected human brain tissue samples were used as input. Nuclei were incubated overnight at 4°C with a primary MITF antibody (Abcam, Cat. #ab12039, mouse; 1:100 dilution). The following day, MeA/MeRev-loaded nano-Tn5 anti-mouse (provided by Protein Science Facility, Karolinska Institutet) was added and incubated for 1 hour at room temperature. After washing, tagmentation was carried out at 37°C for 1 hour. DNA was purified, and a first linear amplification was performed using the i5 primer only (10 cycles). A second tagmentation using commercial Tn5 (Diagenode, C01070010), loaded with MeB/MeRev adaptors, was followed by exponential amplification with both i5 and i7 primers (11 cycles) to generate the final sequencing libraries.

### Processing of MITF CUT&Tag fastq files

Raw reads of MITF-CUT&Tag data were processed with the nf-core/cutandrun pipeline^83,110^. Post-alignment, low-quality reads were filtered out, and bam files were merged and indexed using Samtools (v1.21). Peak calling was performed using MACS2 (v2.2.9.1)^111^ on merged bam files using the parameters: -q 1e-5 -f BAMPE –nomodel –shift -75 –extsize 150 – nolambda. Peaks mapping to mitochondria or random chromosomes were removed. For downstream analyses, only high confidence peaks with p ≤ 1 x 10^-7^ were retained.

### MITF CUT&Tag analysis

MITF CUT&Tag binding sites that overlap with microglia H3K27ac were identified in R using GenomicRanges^89^ and were assigned to genes with annotatePeak using reference transcriptome TxDb.Hsapiens.UCSC.hg38.refGene and annotation database org.Hs.eg.db. Gene ontology enrichment analysis of the genes annotated to MITF binding sites at H3K27ac regions was assessed using enrichGO from clusterProfiler^112^. HOMER *de novo* motif discovery of MITF binding sites was performed using all MITF CUT&Tag peaks as input and automatic background, as described in the ‘transcription factor motif analysis’ section.

MITF signal at microglia AD-associated H3K27ac peaks was assessed using aggregate plot analysis. MITF signal at acetylated regions was determined using computeMatrix reference-point from deeptools^113^ with further processing of the output matrix using plotProfile. Inactive regions were identified as regions 2000bp upstream and downstream of TSSs that do not overlap with H3K27ac peaks.

Genes annotated to AD differential H3K27ac peaks at MITF binding sites in microglia were assessed for enrichment of cell subtype signature genes using Fisher’s exact test. Signature genes for 8 substates were defined using a human-mouse amyloid model by scRNA-seq^25^. Fisher’s exact test (v4.3.1) was performed as described in the ‘cell subtype enrichment analysis’ section.

